# Assessing RNAi feasibility and susceptibility to environmental RNAi in *Trichogramma dendrolimi* (Hymenoptera: Trichogrammatidae)

**DOI:** 10.1101/2023.07.03.547498

**Authors:** Zhichao Yan, Fangyi Li, Aokai Wang, Chengxing Wang, Haiyan Wang, Zeqi Yu, Kepeng Wang, Yihan Wang, Yuanyuan Luo, Yuanxi Li

## Abstract

*Trichogramma*, a genus of egg parasitoid wasps, are widely used as biological control agents and serve as model organisms in parasitoid research. Despite their significance, the understanding of RNA interference (RNAi) in *Trichogramma* remains very limited. In this study, we investigated RNAi-associated genes by bioinformatic approaches and experimentally assessed the feasibility of RNAi and the susceptibility of environmental RNAi in *Trichogramma*. We found that *Trichogramma* genomes contain a complete set of genes in the RNAi pathway and exhibit extensive gene expansion of dsRNase, which may influence RNAi efficiency by degrading dsRNA. We demonstrated successful RNAi through pupal microinjection in *T. dendrolimi* Matsumura, providing a technical approach for future gene functional studies. In addition, we observed no evidence of susceptibility to environmental RNAi in either *T. dendrolimi* adults or larvae, which might be attributed to the extensive expansion of dsRNase. This low environmental RNAi sensitivity in *Trichogramma* could suggest a reduced risk of RNAi-based pest management strategies affecting nontarget *Trichogramma* populations. Overall, this study presents a technical approach for conducting gene functional studies in *Trichogramma* and provides a foundation for evaluating the nontarget effects of RNAi-based pest control strategies on *Trichogramma*.

## Introduction

RNA interference (RNAi) is a conserved gene regulatory mechanism found in eukaryotes (Carthew and Sontheimer 2009; Wilson and Doudna 2013). The underlying mechanism of RNAi mainly involves the generation of small RNA molecules (such as siRNA and miRNA) by Dicer cleavage (Vergani-Junior et al. 2021). These small RNAs subsequently bind to Argonaute proteins to form RNA-induced silencing complexes (RISC), leading to the degradation or translational repression of target mRNA (Iwakawa and Tomari 2022; Meister 2013). RNAi plays a vital role in gene expression regulation and affects almost all biological processes (Wilson and Doudna 2013). In addition to RNAi within individual organisms, there is also accumulating evidence that certain small RNAs can be exchanged between hosts and their pathogens or parasites to induce gene silencing, which is known as "cross-kingdom RNAi" (Asgari 2023; Cai et al. 2018; Wang et al. 2017). For example, miRNAs from the parasitoid wasp *Cotesia vestalis* have been shown to suppress the host ecdysone receptor gene to arrest host development (Wang et al. 2018).

Since its discovery, RNAi has emerged as a powerful tool for gene functional studies (Mohr et al. 2010; Mohr et al. 2014; Perrimon et al. 2010). It also shows significant applications in pest management (Mezzetti et al. 2020; Zhang et al. 2013). Two prominent RNAi-based strategies are host-induced gene silencing (HIGS) and spray-induced gene silencing (SIGS) (Mezzetti et al. 2020). HIGS is the use of transgenic RNAi crops to silence gene expression in pests, while SIGS involves the direct application of dsRNA pesticides. Despite the conservation of the core RNAi mechanism in insects, there is substantial variation in the efficiency of RNAi (Cooper et al. 2019). Several factors can influence RNAi efficiency across insect species, including the variation in dsRNA degradation, the efficiency of cellular uptake and endosomal escape, and the systemic spreading of RNAi effects (Cooper et al. 2019). A well-studied factor is the presence and abundance of dsRNases, which can degrade dsRNAs and thereby reduce RNAi efficiency (Cooper et al. 2019). It has been reported in several insects that knockdown of dsRNases can improve RNAi efficiency (Ahmad et al. 2023; Fan et al. 2021; Peng et al. 2020; Peng et al. 2021; Sharma et al. 2021; Song et al. 2019; Zhang et al. 2022).

*Trichogramma* are egg parasitoid wasps and one of the most widely used biological control agents, targeting various lepidopteran pests in agriculture and forestry (Knutson 1998; Smith 1996; Zang et al. 2021). They also serve as model organisms in parasitoid research and can be considered “the *Drosophila* in the parasitoid world” (Smith 1996). However, due to their tiny size (∼0.5 mm), the understanding of RNAi in *Trichogramma* remains very limited, largely hindering gene functional studies within the species (Zang et al. 2021). Moreover, *Trichogramma* adults can be directly exposed to RNAi molecules through the spraying of dsRNA pesticides or the consumption of pollen and nectar from transgenic RNAi crops (Christiaens et al. 2022). Indirect exposure could also occur in *Trichogramma* larvae during development inside hosts, which are targeted by HIGS or SIGS (Christiaens et al. 2022). Therefore, it is critical to assess RNAi feasibility and the sensitivity of environmental RNAi in *Trichogramma* to both advance *Trichogramma* research and evaluate nontarget risks on *Trichogramma* for RNAi-based pest management strategies.

In this study, we analyzed genes involved in the RNAi pathway and dsRNase genes in *Trichogramma* genomes. We also established an RNAi method by pupal microinjection and assessed the environmental RNAi sensitivity in *T. dendrolimi*.

## Materials and Methods

### Insect rearing

*Trichogramma dendrolimi* were reared in *Drosophila* tubes in an incubator with a temperature of 25±1°C, a relative humidity of 70±5%, and a light cycle of L:D = 14:10 h (Yan et al. 2022). *Corcyra cephalonica* eggs were used as hosts. After adult wasps emerge, they were fed with 10% sucrose solution to extend their lifespan.

### Identification and analysis of RNAi-related genes

To identify genes involved in the RNAi pathway in *Trichogramma* genomes, BLASTP v2.12.0+ was performed against the genomes of *T. dendrolimi* and *T. pretiosum* (Riley) (Lindsey et al. 2018) using query sequences from *Drosophila melanogaster* Meigen, *Tribolium castaneum* Herbst, *Bombyx mori* L., and *Apis mellifera* L. (for details, see Supplementary Text). Protein domains were predicted using the Pfam (http://pfam-legacy.xfam.org/) (Mistry et al. 2021) and Interpro (http://www.ebi.ac.uk/interpro/) (Paysan-Lafosse et al. 2023) online services and visualized using TBTools v1.098769 (Chen et al. 2020). Protein sequences were aligned using Muscle (Edgar 2004), which was built in MEGA v10.2.3 (Kumar et al. 2018). Maximum likelihood phylogenies were constructed using MEGA v10.2.3 with bootstrap values of 1000 (Kumar et al. 2018). For dsRNase identification, domain PF01223.26 (Endonuclease_NS) was scanned in insect genomes using HMMER v3.3.2 with an e-value of 1e^-3^ (Mistry et al. 2013). Alignments were performed using Muscle v4.0.0.0 (Edgar 2004) and trimmed by trimAl v1.4 (Capella-Gutiérrez et al. 2009). The phylogenetic tree was constructed using IQTree v4.2.3 (Minh et al. 2020) and visualized using iTOL v6 (Letunic and Bork 2021). Heatmaps of gene expression were drawn using TBtools v1.098769 (Chen et al. 2020).

### dsRNA synthesis

*Trichogramma dendrolimi* wasps, 5 to 6 days after eclosion, were collected for RNA extraction. Total RNA was extracted using TRIzol reagent (Invitrogen, USA) and reverse-transcribed into cDNA using a HiScript III RT SuperMix for qPCR (+gDNA wiper) kit (Vazyme, Nanjing, China). Primers were designed using Primer Premier 5 (Premier Biosoft Inc., USA) by setting product lengths of 300–500 bp, primer lengths of 21±3 bp, and primer GC% of 40%–60%. Primers are listed in Table S1. The T7 promoter sequence 5′-TAATACGACTCACTATA-3′ was then manually added to the 5’ end of all primers. Templates for dsRNA synthesis were amplified using Rapid Taq Master Mix (Vazyme, Nanjing, China) and purified using the AxyPrep DNA Gel Recovery Kit (Axygen, USA). dsRNA was synthesized using the T7 High Yield RNA Transcription Kit (Vazyme, Nanjing, China).

### Delivery of dsRNA

For the pupal stage, dsRNA was delivered by microinjection. *Trichogramma dendrolimi* prepupae were carefully extracted from parasitized *C. cephalonica* eggs after four days of parasitism and arranged on a glass slide. dsRNA (∼10 μg/μl) was microinjected into the abdomen of the prepupae using a FemtoJet® 4i (Eppendorf, Germany), with the injected droplet size being comparable to the wasp’s compound eye. Injected prepupae were sealed in artificial egg cards made of polyethylene film and placed in a Petri dish accompanied by a moistened cotton ball for humidity control. The knockdown of target genes was evaluated by RT‒qPCR 48 h post-injection.

For the adult stage, dsRNA was delivered through *ad lib* feeding of *T. dendrolimi* wasps. A 10% sucrose solution containing dsRNA (1 μg/μl) and erioglaucine blue colorant (0.05 mg/ml, Sigma) was offered to newly emerged wasps via capillary glass tubes with a 0.5 mm inner diameter. When using BAPtofect™-25 (for insect only) (Phoreus Biotech, USA) to coat dsRNA (Avila et al. 2018), it was mixed with dsRNA at a 20:1 mass ratio according to the manufacturer’s instructions. The dsRNA-containing capillary tubes were replaced every 24 h. The knockdown of target genes was evaluated by RT‒qPCR 72 h after *ad lib* feeding.

For the larval stage, dsRNA was delivered using artificial hosts. Artificial hosts were prepared as previously described (Lü et al. 2017), with the addition of 1 μg/μl dsRNA. When BAPtofect™-25 was used to coat the dsRNA (Avila et al. 2018), it was combined with dsRNA in a 20:1 mass ratio according to the manufacturer’s instructions. Mated wasps that had newly eclosed within 24 h were used to parasitize artificial hosts for 4 h. Each parasitized artificial host was carefully opened to add 1 μl of 3300 ng/μl dsRNA solution at 24 h post-parasitism, and subsequently resealed for continued incubation. The process of parasitism and incubation of artificial hosts was conducted in an incubator (27±1℃, RH 80±5%, L:D = 14:10 h). The knockdown of target genes was evaluated by RT‒qPCR 72 hours post-parasitism.

### RT‒qPCR

Primers for RT‒qPCR were designed by Primer Premier 5 (Premier Biosoft Inc., USA) to avoid overlapping with the regions of synthesized dsRNA(Onchuru and Kaltenpoth 2019). Primers are listed in Table S2. Total RNA was extracted using TRIzol reagent and reverse transcribed using the HiScript III RT SuperMix for qPCR (+gDNA wiper) kit (Vazyme, Nanjing, China). qPCRs were conducted on a QuantStudio™ 5 instrument (Applied Biosystems, USA) using ChamQ Universal SYBR qPCR Master Mix (Vazyme, Nanjing, China). The qPCR cycling conditions were as follows: initial denaturation at 95°C for 30 s, followed by 40 cycles of 10 s at 95°C and 30 s at 60°C, and a final step consisting of 15 s at 95°C, 60 s at 60°C, and 15 s at 95°C. The actin gene was used as an internal reference gene, and the relative expression levels of the target genes were calculated using the 2^-ΔΔCT^ method.

### Eclosion rate and survival curve analysis

Beginning on the eighth day post-parasitism, the emergence of injected *T. dendrolimi* pupae was monitored at 24-hour intervals for a duration of 5 days. Typically, eclosion begins between 9 and 10 days post-parasitism. Newly emerged adult wasps were transferred to new *Drosophila* tubes and fed a 10% sucrose solution. Subsequently, the survival of the wasps was recorded at 12 h intervals over a consecutive 5-day period.

### Statistical analyses

Statistical analyses were performed using GraphPad Prism 9.0 (GraphPad Software, San Diego, USA). Eclosion rates were first subjected to arcsine square root transformation. Two-tailed Student’s t test was used to analyze differences between two samples, and one-way ANOVA was performed for multiple group comparisons. Survival curve data were analyzed using the log-rank test.

## Results

### *Trichogramma* genomes contain a complete set of genes in the RNAi pathway and show an expansion of dsRNase genes

Using query proteins of the RNAi pathway from *Drosophila melanogaster*, *Tribolium castaneum*, *Bombyx mori*, and *Apis mellifera*, we identified 2 Dicer, 1 Drosha, 3 Argonaute, 1 Lops, 1 Pasha, 1 R2D2, and 2 Sid-1 genes in the genomes of both *T. dendrolimi* and *T. pretiosum* (Fig. 1a, S1-S8; Table 1). Domain analyses showed that these genes contained conserved protein domains and domain arrangements (Fig. S1, 3, 5 & 7). Notably, the phylogeny of Argonaute suggested a recent gene duplication of Ago-1 after the divergence of *Trichogramma* from *Apis* (Fig. S4). For Sid-1, both the *T. dendrolimi* and *T. pretiosum* genomes contain two gene copies, while most other insects contain only one copy (Fig. S8). More details are provided in the supplementary text. These results demonstrate the presence of a complete set of genes encoding for the RNAi pathway in the genomes of both *T. dendrolimi* and *T. pretiosum*.

**Figure 1.**
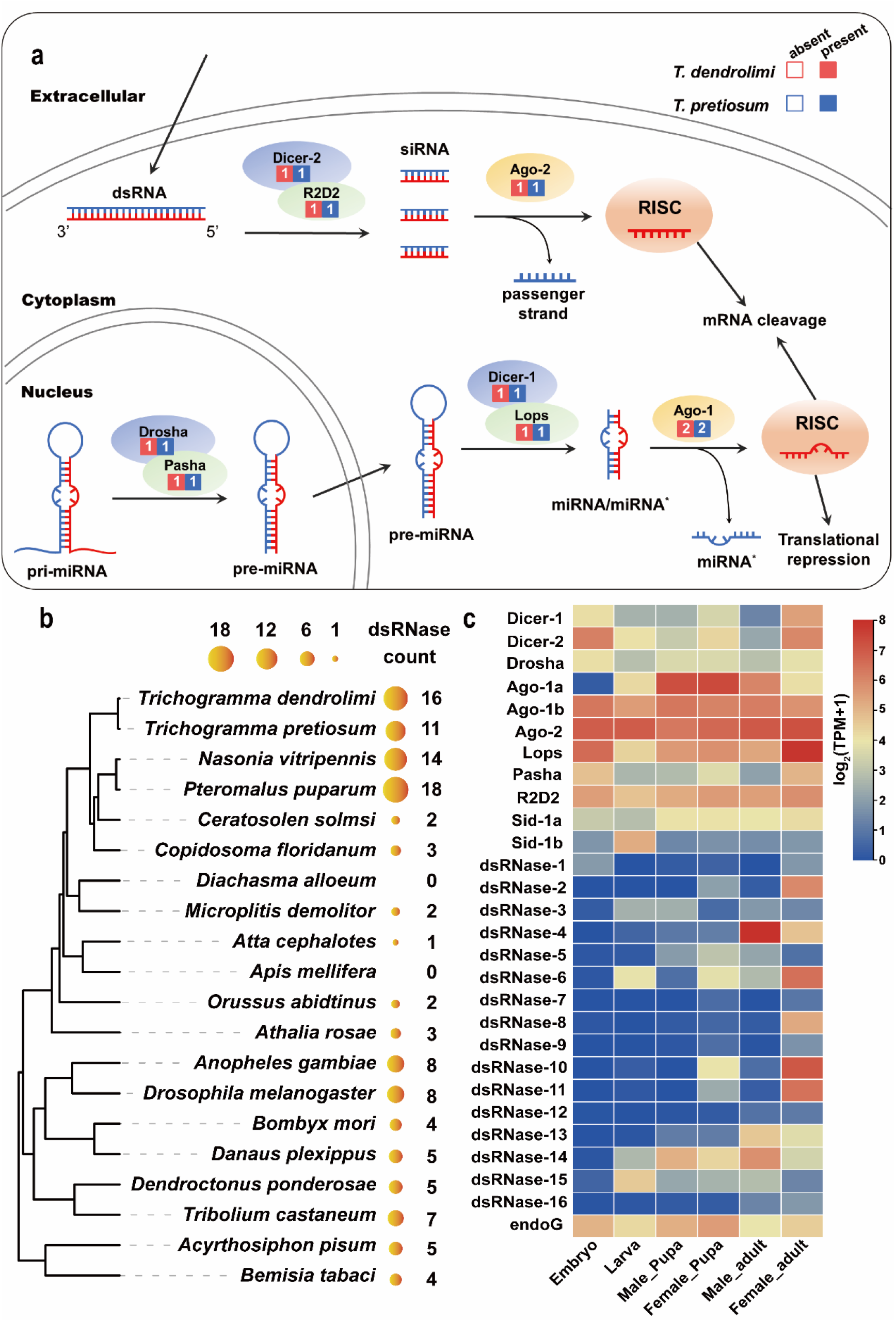
*Trichogramma* genomes contain the complete set of genes in the RNAi pathway and show extensive expansion of dsRNase genes. (a) Identification of genes involved in the RNAi pathway in *T. dendrolimi* and *T. pretiosum*. The numbers in the boxes represent the number of identified genes. * indicates the antisense strand. (b) Number of identified dsRNase genes in insect genomes. (c) Expression levels of RNAi-related genes across different developmental stages in *T. dendrolimi*.

**Table 1.** Characteristics of RNAi pathway-related genes in *Trichogramma dendrolimi* and *T. pretiosum*.

| Gene | Length (aa.) | IP | MW (kDa) | Accession number |
| --- | --- | --- | --- | --- |
| *T. dendrolimi* |  |  |  |  |
| Dicer-1 | 1899 | 5.84 | 217.70 | Tden08220.t1 |
| Dicer-2 | 1654 | 6.19 | 190.40 | Tden14400.t1 |
| Drosha | 1428 | 9.15 | 164.86 | Tden01949.t1 |
| Ago-1a | 896 | 9.23 | 100.82 | Tden05973.t1 |
| Ago-1b | 950 | 9.14 | 106.47 | Tden09851.t1 |
| Ago-2 | 1189 | 9.69 | 132.86 | Tden08262.t24 |
| Lops | 379 | 7.63 | 42.15 | Tden13462.t1 |
| Pasha | 634 | 6.35 | 71.97 | Tden12859.t1 |
| R2D2 | 371 | 5.58 | 41.79 | Tden00172.t1 |
| Sid-1a | 793 | 6.68 | 91.07 | Tden08522.t1 |
| Sid-1b | 595 | 6.20 | 67.50 | Tden08523.t2 |
| *T. pretiosum* |  |  |  |  |
| Dicer-1 | 1899 | 5.89 | 217.59 | XP_014235184.1 |
| Dicer-2 | 1654 | 6.05 | 189.86 | XP_014238228.1 |
| Drosha | 1429 | 9.08 | 164.89 | XP_014223505.1 |
| Ago-1a | 896 | 9.23 | 100.82 | XP_014225266.1 |
| Ago-1b | 951 | 9.14 | 106.55 | XP_014228304.1 |
| Ago-2 | 1087 | 9.54 | 122.20 | XP_023316532.1 |
| Lops | 332 | 6.76 | 36.82 | XP_014228041.1 |
| Pasha | 652 | 6.43 | 74.12 | XP_023316230.1 |
| R2D2 | 347 | 5.42 | 39.08 | XP_014221168.2 |
| Sid-1a | 793 | 7.03 | 91.08 | XP_014230770.1 |
| Sid-1b | 820 | 6.65 | 93.70 | XP_014230830.1 |

aa. : amino acids; IP: isoelectric point; MW: molecular weight.

As dsRNases can influence insect RNAi efficiency by degrading dsRNA, we further investigated the presence of dsRNase genes in several representative insects. We identified 16 and 11 dsRNase genes in the genomes of *T. dendrolimi* and *T. pretiosum*, respectively (Fig. 1b; Table S3). The count of dsRNase was 18 in *Pteromalus puparum* and 14 in *Nasonia vitripennis*, but the count ranged from 0 to 3 in the other surveyed hymenopteran insects (Fig. 1b). Phylogenetic analysis suggests that the expansion of dsRNase in *Trichogramma* is mainly Chalcidoidea specific, with some unique gene expansions observed in *Trichogramma* (Fig. 2).

**Figure 2.**
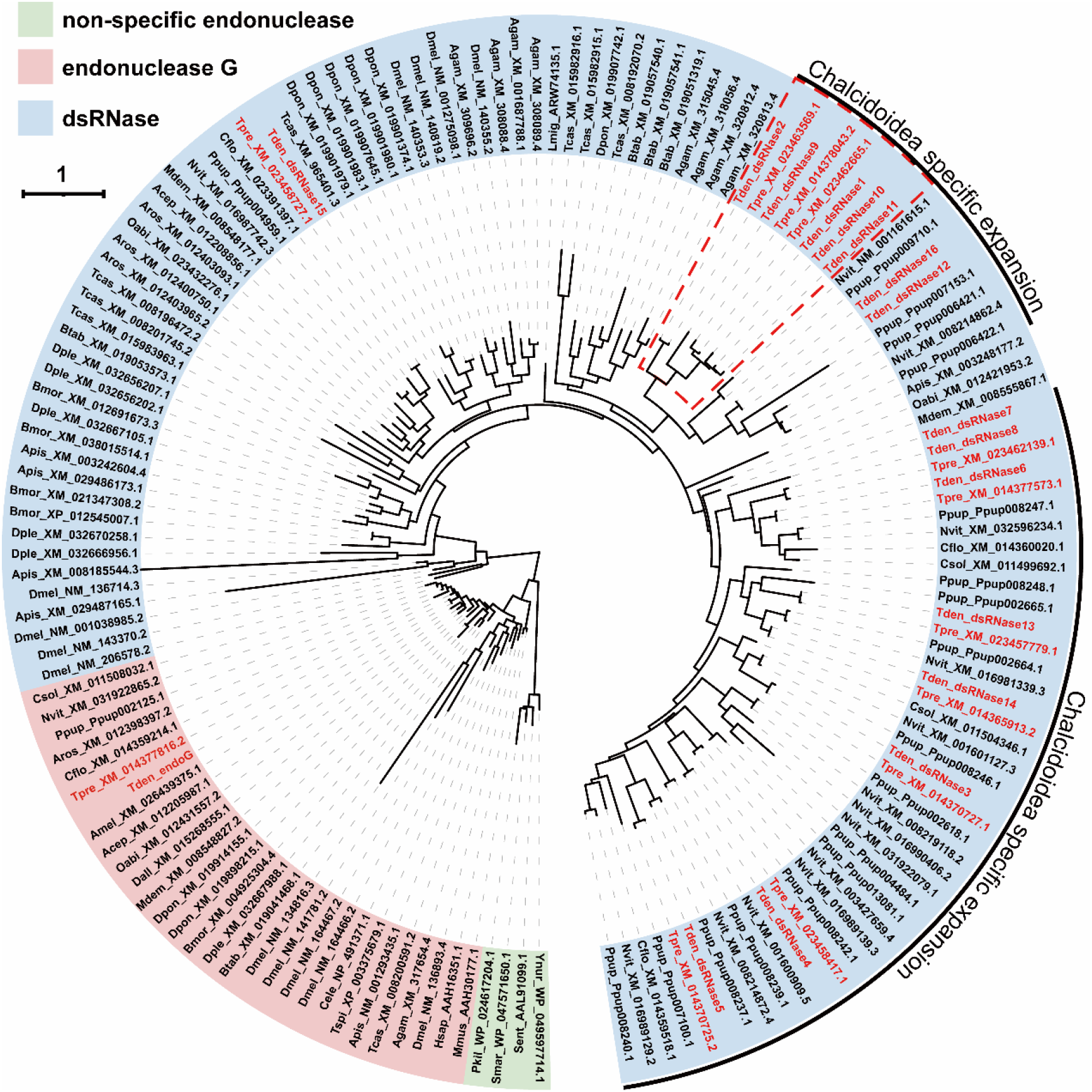
The maximum-likelihood phylogenetic tree of dsRNase. Genes from *T. dendrolimi* and *T. pretiosum* are labeled in red. The dashed red box indicates *Trichogramma*-specific expansion of dsRNase. Agam: *Anopheles gambiae*; Amel: *Apis mellifera*; Acep: *Atta cephalotes*; Apis: *Acyrthosiphon pisum*; Aros: *Athalia rosae*; Bmor: *Bombyx mori*; Btab: *Bemisia tabaci*; Cflo: *Copidosoma floridanum*; Csol: *Ceratosolen solmsi*; Dall: *Diachasma alloeum*; Dmel: *Drosophila melanogaster*; Dple: *Danaus plexippus*; Dpon: *Dendroctonus ponderosae*; Mdem: *Microplitis demolitor*; Nvit: *Nasonia vitripennis*; Oabi: *Orussus abidtinus*; Pkil: *Pseudomonas kilonensis*; Ppup: *Pteromalus puparum*; Sent: *Salmonella enterica*; Smar: *Serratia marcescens*; Tcas: *Tribolium castaneum*; Tden: *Trichogramma dendrolimi*; Tpre: *Trichogramma pretiosum*; Ynur: *Yersinia nurmii*.

Using RNA-seq data, we examined the gene expression of the RNAi pathway across different developmental stages. The genes in the RNAi pathway generally showed high expression levels across all stages, except for Ago-1a, which showed low expression in embryos (Fig. 1c). In contrast, dsRNase expression levels appeared to be stage specific (Fig. 1c). For example, dsRNase-4 was differentially highly expressed in male adults, while dsRNase-2, 8, 10, and 11 were differentially highly expressed in female adults.

### Efficient RNAi can be achieved by injecting dsRNA into *T. dendrolimi* pupae

For RNAi experiments, we selected three target genes, namely, chitinase 10 (CHT10), v-ATPase A (VAA), and v-ATPase B (VAB), which have been reported in other insects to affect their survival or molting (Table 2) (Joga et al. 2016; Mehlhorn et al. 2021). By microinjection of dsRNA into *T. dendrolimi* prepupae, we successfully achieved a significant knockdown in the expression levels of all three genes (Fig. 3). CHT10 showed a reduction of ∼70% in expression (Fig. 3c; Student’s t test; *t* = 8.29, *p* = 0.0012); VAA showed a decrease of ∼28% (Fig. 3f; Student’s t test; *t* = 3.31, *p* = 0.030); and VAB displayed a reduction of ∼30% (Fig. 3i; Student’s t test; *t* =11.03, *p* = 0.0004). These results indicate that injection of dsRNA into *Trichogramma* pupae can effectively suppress the expression of target genes.

**Figure 3.**
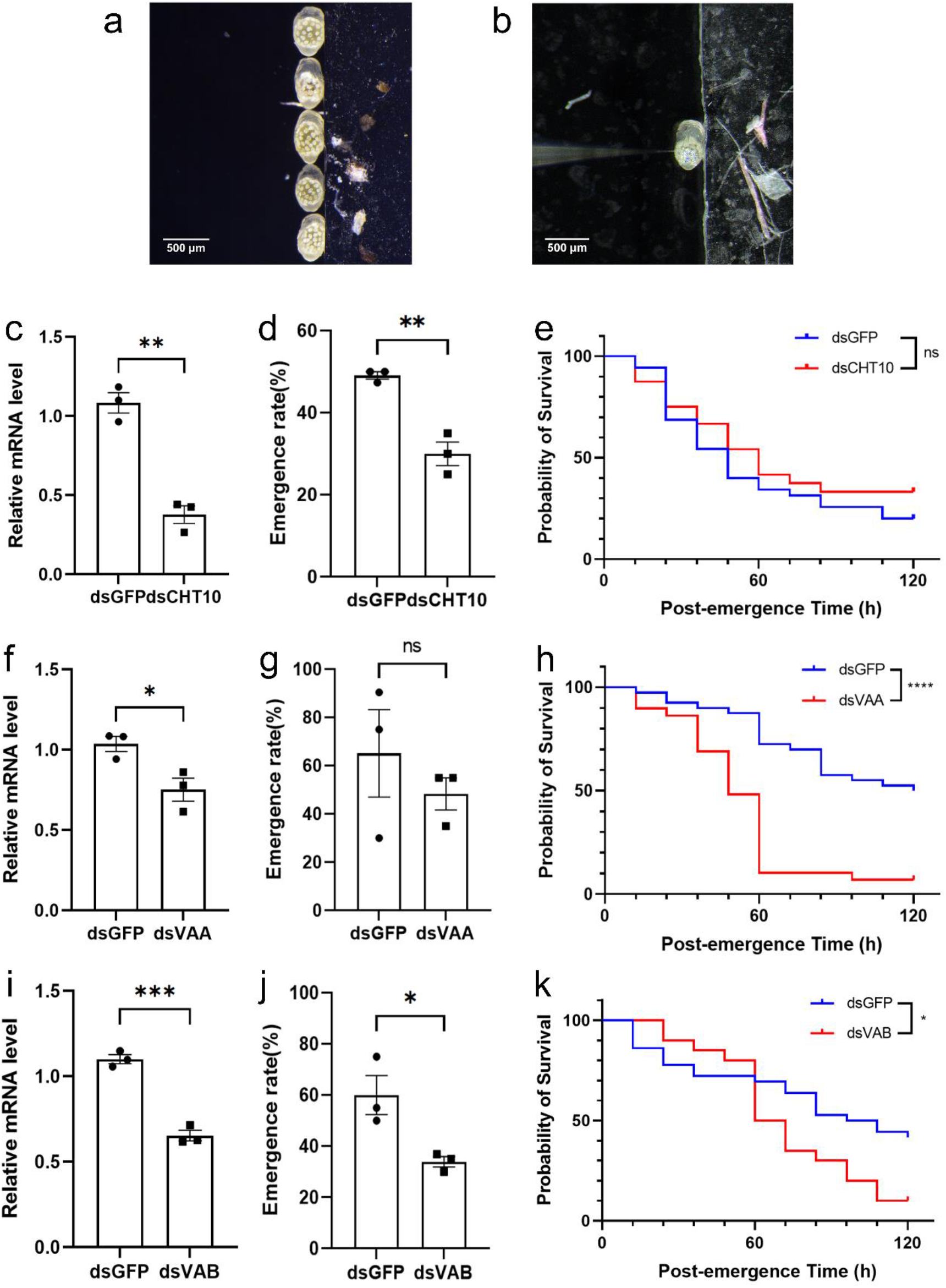
Microinjection of dsRNA into *T. dendrolimi* pupae results in knockdown of target genes and phenotypic changes. (a) *Trichogramma dendrolimi* prepupae 4 d after parasitism. (b) Microinjection of dsRNA into *T. dendrolimi* prepupae. (c, f, i) Relative expression levels of (c) CHT10, (f) VAA and (i) VAB in *T. dendrolimi* pupae 48 h after microinjection of dsRNA. (d, g, j) Eclosion rate of *T. dendrolimi* pupae after microinjection of dsRNA of (d) CHT10, (g) VAA and (j) VAB. (e, h, k) Survival curve of emerged *T. dendrolimi* adults after microinjection of dsRNA of (e) CHT10, (h) VAA and (k) VAB. Data are displayed as the mean ± SEM, with n=3. Ns: no significant difference; * *p* < 0.05; **: *p* < 0.01; ***: *p* < 0.001.

**Table 2.**
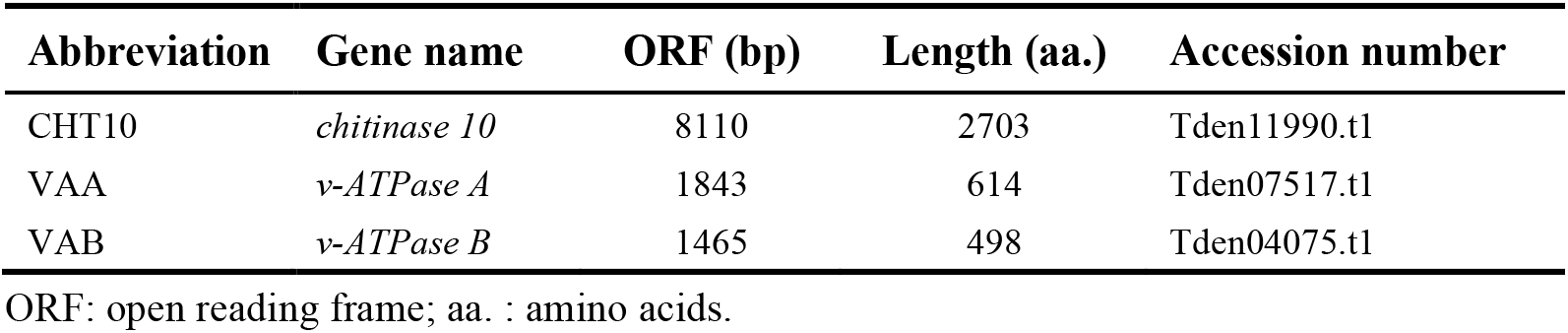
Target genes for microinjection RNAi in *T. dendrolimi*.

Moreover, RNAi of these target genes affected *T. dendrolimi* eclosion and survival. After CHT10 RNAi, the eclosion rate of *T. dendrolimi* was significantly reduced (Fig. 3d; Student’s t test; *t* = 6.1820, *p* = 0.000842), but the adult survival curve showed no significant difference compared to the control group (Fig. 33; log-rank test; *p* = 0.3252). After VAA RNAi, the eclosion rate showed no significant difference from the control (Fig. 3g; Student’s t test; *t* = 0.8707, *p* = 0.4331), but the adult survival curve was significantly reduced (Fig. 3h; log-rank test; *p* < 0.0001). After VAB RNAi, both the eclosion rate (Fig. 3j; Student’s t test; *t* = 3.22, *p* = 0.032) and adult survival curve were significantly reduced (Fig. 3k; log-rank test; *p* = 0.030). These phenotypic results following RNAi further support that the knockdown of these target genes is efficient.

### *Trichgoramma dendrolimi* larvae and adults are insensitive to environmental RNAi

To assess the susceptibility of *T. dendrolimi* adult wasps to environmental RNAi, we fed wasps with dsRNA (Fig. a & b). To improve the delivery efficiency of dsRNA, we also applied BAPtofect™-25 nanocarriers to form branched amphipathic peptide capsules for encapsulating dsRNA as previously described in other insects (Avila et al. 2018). Following 3 days of dsRNA feeding, no significant changes were detected in the expression levels of target genes (Fig. 4c-e; one-way ANOVA; all comparisons: *p* > 0.05), regardless of whether the nanocarrier BAPtofect™-25 was used for dsRNA encapsulation. Furthermore, we examined eight additional genes, primarily chosen based on their widespread expression in different stages or our interest in venom (Table S4). After 3 days of dsRNA feeding, no significant knockdown was detected (Fig. S9; Student’s t test; all comparisons: *p* > 0.05).

**Figure 4.**
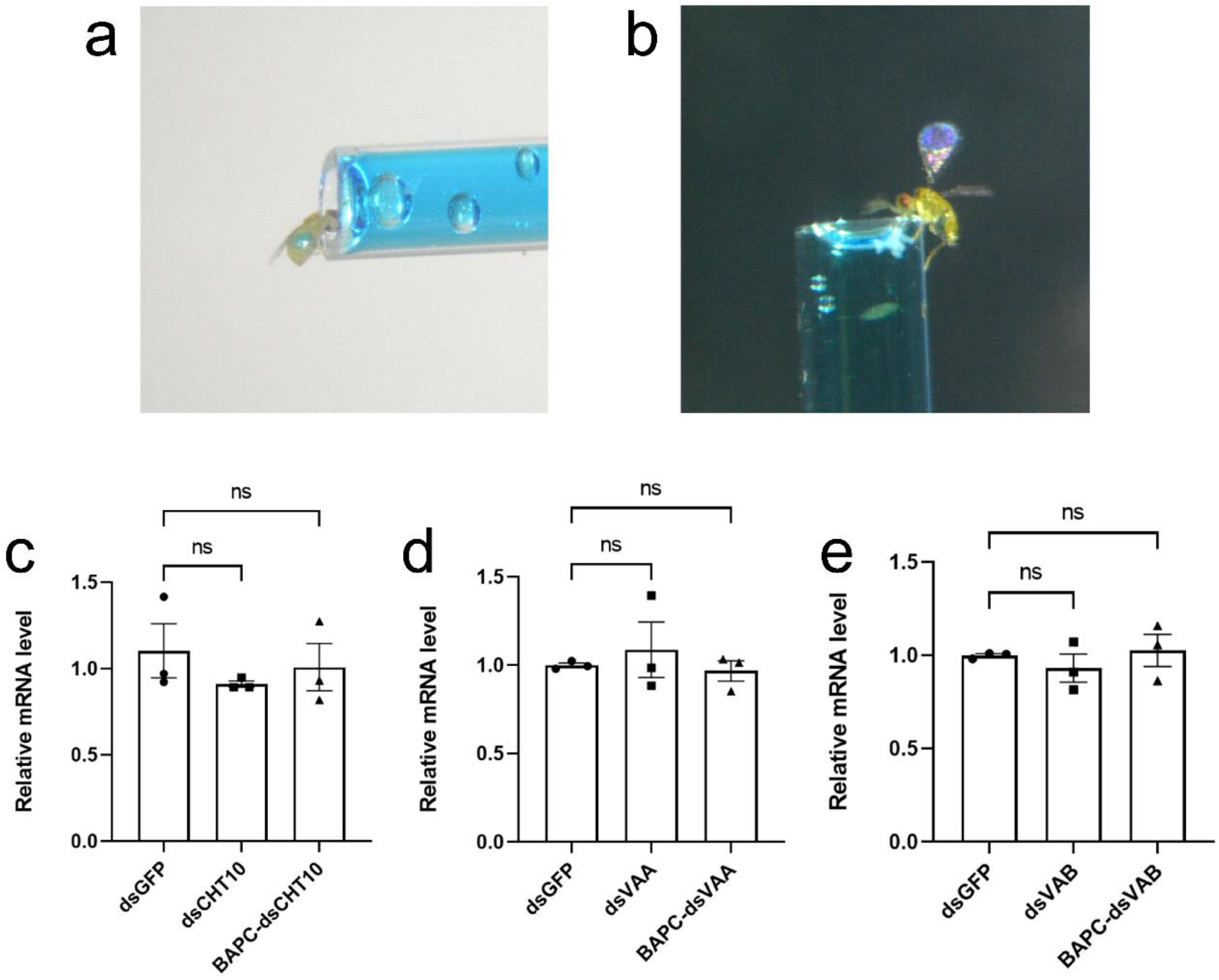
Lack of knockdown effects in *T. dendrolimi* adults following 3 days of continuous dsRNA feeding. (a, b) *Trichogramma dendrolimi* adults consuming 10% sucrose solution with dsRNA. The inner diameters of capillary glass tubes are 0.5 mm. (c) Relative expression levels of (c) CHT10, (d) VAA, and (e) VAB in *T. dendrolimi* adults after feeding on dsRNA with or without BAPC encapsulation. GFP dsRNA served as a negative control. BAPC: branched amphipathic peptide capsule. Data are displayed as the mean ± SEM, with n=3. Ns: no significant difference.

To assess the susceptibility of *T. dendrolimi* larvae to environmental RNAi, we provided artificial hosts containing dsRNA for *Trichogramma* to parasitize. *Trichogramma dendrolimi* can successfully lay eggs and develop within these artificial hosts (Fig. 5a, b). BAPtofect™-25 was also applied for dsRNA encapsulation. No significant changes were observed in the expression levels of the target genes CHT10, VAA, and VAB, regardless of whether BAPtofect™-25 was used (Fig. 4c-h, t test, *p* > 0.05). Similarly, we also examined 3 additional genes and found no knockdown of target genes (Fig. S10; Student’s t test; all comparisons: *p* > 0.05).

**Figure 5.**
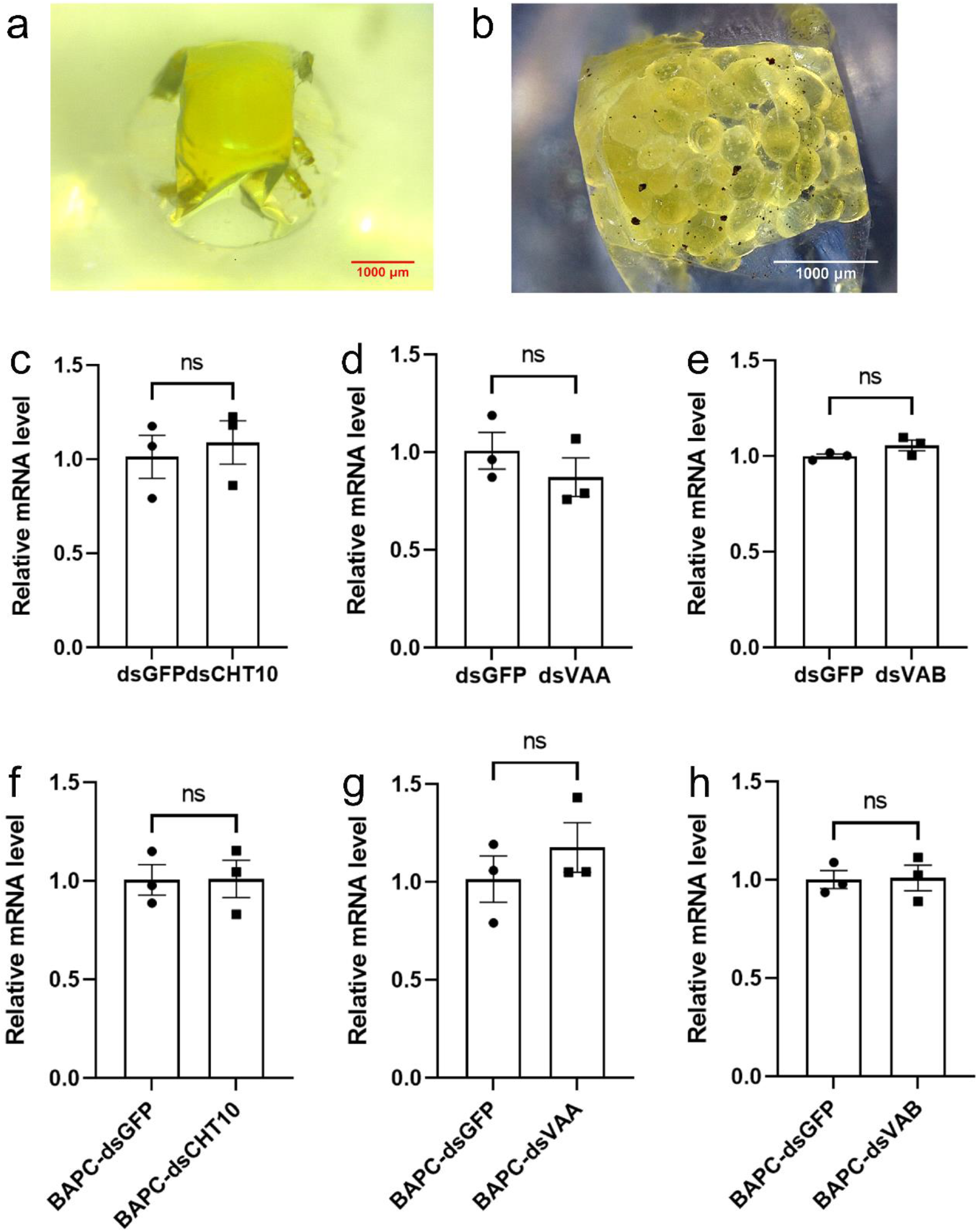
Lack of knockdown effects in *T. dendrolimi* larvae parasitized in artificial hosts containing dsRNA. (a) *Trichogramma dendrolimi* adults parasitizing artificial hosts. (b) *Trichogramma dendrolimi* larvae 3 days after developing in artificial hosts. (c-e) Relative expression levels of (c) CHT10, (d) VAA, and (e) VAB in *T. dendrolimi* larvae 3 days after parasitism in artificial hosts containing dsRNA. (f-h) Relative expression levels of (f) CHT10, (g) VAA, and (h) VAB in *T. dendrolimi* larvae 3 days after parasitism in artificial hosts containing dsRNA with BAPC capsulation. GFP dsRNA served as a negative control. BAPC: branched amphipathic peptide capsule. Data are displayed as the mean ± SEM, with n=3. Ns: no significant difference.

## Discussion

In this study, we investigated RNAi-associated genes by bioinformatic approaches and experimentally assessed the feasibility of RNAi and the susceptibility of environmental RNAi in *Trichogramma*. We found that both *T. dendrolimi* and *T. pretiosum* contain a complete set of genes encoding for the RNAi pathway. These genes showed high expression levels across different developmental stages. To verify the feasibility of RNAi in *Trichogramma*, we microinjected dsRNA into *T. dendrolimi* pupae. Both RT‒ qPCR and phenotypic results confirmed successful knockdown of target genes. This injection-based RNAi method demonstrates the feasibility of RNAi in *Trichogramma*, providing a pathway for future research on *Trichogramma* gene functions.

In addition, we are particularly interested in the susceptibility of environmental RNAi in *Trichogramma* for several reasons. First, compared to microinjection, RNAi by feeding or soaking can significantly increase the knockdown throughput, especially for tiny *Trichogramma* wasps. Second, susceptibility evaluation of *Trichogramma* to environmental RNAi can provide clues for the possibility of cross-kingdom RNAi between *Trichogramma* and their hosts. Third, it is also critical for the risk assessment of HIGS and SIGS on *Trichogramma*. To this end, we assessed the environmental RNAi susceptibility of *T. dendrolimi* adults and larvae. In neither adults nor larvae, no significant knockdown effects were observed even with the use of the nanocarrier BAPtofect™-25 for dsRNA encapsulation (Avila et al. 2018). These results indicate that *T. dendrolimi* has no or low sensitivity to environmental RNAi.

Correspondingly, we observed extensive gene expansion of dsRNase in *Trichogramma*, with some of these expansions being *Trichogramma* specific. Given that dsRNase can degrade dsRNA and thereby reduce RNAi efficiency, the extensive gene expansion of dsRNase in *Trichogramm*a may explain its low sensitivity to environmental RNAi. For dsRNases, the expression widths across developmental stages are generally narrower compared to the essential mitochondrial gene endonuclease G (endoG) and the core genes involved in the RNAi pathway. The expression of several dsRNases appears to be stage specific. However, due to the lack of tissue expression data, it remains unclear whether these dsRNases are enriched at the interface between *Trichogramma* and their hosts during parasitization or host feeding, such as venom, larval saliva, and adult saliva. One possible explanation for the expansion of dsRNase genes is to protect *Trichogramma* from attacks by host small RNAs. Further investigations are needed to test these hypotheses.

Moreover, *Trichogramma* are commonly used as nontarget indicator organisms for the risk assessment of genetically modified crops and pesticides (O’Callaghan et al. 2005; Rakes et al. 2021). In the context of RNAi-based pest control strategies, *Trichogramma* can directly or indirectly come into contact with insecticidal RNAi molecules derived from HIGS or SIGS approaches. Regarding the sensitivity to environmental RNAi, the findings in *Trichogramma* differ from a previous report on the egg parasitoid *Telenomus podisi*, which is sensitive to dietary intake of dsRNA (Castellanos et al. 2022). In contrast, *T. dendrolimi* is insensitive to dietary dsRNA in both adult and larval stages. The low environmental RNAi sensitivity of *Trichogramma* may suggest a reduced risk of HIGS and SIGS on *Trichogramma* populations. However, it is important to note that genetically modified plants can be engineered to direct the accumulation of dsRNA in plant chloroplasts, thereby enhancing dsRNA resistance against digestion in insect guts. Similarly, dsRNA pesticides can utilize various nanocarriers for dsRNA encapsulation. Further investigations are required to evaluate each specific case.

In summary, we found that *Trichogramma* contains a complete set of RNAi pathway genes and shows gene expansion of dsRNase. We demonstrate the feasibility of *Trichgoramma* RNAi through pupal microinjection. We also show that the sensitivity of environmental RNAi is low in *Trichogramma* adults and larvae. These results offer a technical approach for conducting gene functional studies in *Trichogramma* and provide a foundation for evaluating the nontarget effects of RNAi-based pest control strategies on *Trichogramma*.

## Authors’ contributions

ZCY and YXL conceived and designed the research; FYL and CXW conducted microinjection RNAi; ZCY, FYL, AKW performed bioinformatic analyses; FYL, HYW, ZQY, KPW, YHW and YYL performed environmental RNAi; ZCY and YXL interpreted the results; ZCY wrote the first draft of the manuscript; YXL commented on and revised the manuscript; All authors read and approved the final manuscript.

## Acknowledgments

This study was supported by the Natural Science Foundation of Hainan Province (Grant no. 323QN262), the National Natural Science Foundation of China (Grant no. 31701843), the Jiangsu Agriculture Science and Technology Innovation Fund (Grant No. CX(22)3012), the “Shuangchuang Doctor” Foundation of Jiangsu Province (Grant No. 202030472), and the Nanjing Agricultural University startup fund (Grant No. 804018). Bioinformatic analyses were supported by the high-performance computing platform of the Bioinformatics Center, Nanjing Agricultural University.

## References

Ahmad S, Jamil M, Jaworski CC, Luo Y (2023) Double-stranded RNA degrading nuclease affects RNAi efficiency in the melon fly, *Zeugodacus cucurbitae*. J Pest Sci: https://doi.org/10.1007/s10340-023-01637-1.

Asgari S (2023) Cross-kingdom RNAi to enhance the efficacy of insect pathogens. Trends Parasitol 39 (1): 4–6.

Avila LA, et al. (2018) Delivery of lethal dsRNAs in insect diets by branched amphiphilic peptide capsules. J Control Release 273: 139–46.

Cai Q, He B, Kogel KH, Jin H (2018) Cross-kingdom RNA trafficking and environmental RNAi-nature’s blueprint for modern crop protection strategies. Curr Opin Microbiol 46: 58–64.

Capella-Gutiérrez S, Silla-Martínez JM, Gabaldón T (2009) trimAl: a tool for automated alignment trimming in large-scale phylogenetic analyses. Bioinformatics 25 (15): 1972–3.

Carthew RW, Sontheimer EJ (2009) Origins and mechanisms of miRNAs and siRNAs. Cell 136 (4): 642–55.

Castellanos NL, Smagghe G, Taning CNT, Oliveira EE, Christiaens O (2022) Risk assessment of RNAi-based pesticides to non-target organisms: evaluating the effects of sequence similarity in the parasitoid wasp Telenomus podisi. Sci Total Environ 832: 154746.

Chen C, Chen H, Zhang Y, Thomas HR, Frank MH, He Y, Xia R (2020) TBtools: an integrative toolkit developed for interactive analyses of big biological data. Mol Plant 13 (8): 1194–202.

Christiaens O, Sweet J, Dzhambazova T, Urru I, Smagghe G, Kostov K, Arpaia S (2022) Implementation of RNAi-based arthropod pest control: environmental risks, potential for resistance and regulatory considerations. J Pest Sci 95 (1): 1–15.

Cooper AM, Silver K, Zhang J, Park Y, Zhu KY (2019) Molecular mechanisms influencing efficiency of RNA interference in insects. Pest Manag Sci 75 (1): 18–28.

Edgar RC (2004) MUSCLE: a multiple sequence alignment method with reduced time and space complexity. BMC Bioinformatics 5: 113.

Fan YH, et al. (2021) A dsRNA-degrading nuclease (dsRNase2) limits RNAi efficiency in the Asian corn borer (*Ostrinia furnacalis*). Insect Sci 28 (6): 1677–89.

Iwakawa HO, Tomari Y (2022) Life of RISC: formation, action, and degradation of RNA-induced silencing complex. Mol Cell 82 (1): 30–43.

Joga MR, Zotti MJ, Smagghe G, Christiaens O (2016) RNAi efficiency, systemic properties, and novel delivery methods for pest insect control: what we know so far. Front Physiol 7: 553.

Knutson A (1998) The *Trichogramma* manual. Texas Agricultural Extension Service: USA.

Kumar S, Stecher G, Li M, Knyaz C, Tamura K (2018) MEGA X: molecular evolutionary genetics analysis across computing platforms. Mol Biol Evol 35 (6): 1547–49.

Letunic I, Bork P (2021) Interactive Tree of Life (iTOL) v5: an online tool for phylogenetic tree display and annotation. Nucleic Acids Res 49 (W1): W293–W96.

Lindsey ARI, et al. (2018) Comparative genomics of the miniature wasp and pest control agent *Trichogramma pretiosum*. BMC Biol 16 (1): 54.

Lü X, Han S, Li Z, Li L (2017) Biological characters of *Trichogramma dendrolimi* (Hymenoptera: Trichogrammatidae) reared in vitro versus in vivo for thirty generations. Sci Rep 7 (1): 17928.

Mehlhorn S, Hunnekuhl VS, Geibel S, Nauen R, Bucher G (2021) Establishing RNAi for basic research and pest control and identification of the most efficient target genes for pest control: a brief guide. Front Zool 18 (1): 60.

Meister G (2013) Argonaute proteins: functional insights and emerging roles. Nat Rev Genet 14 (7): 447–59.

Mezzetti B, et al. (2020) RNAi: what is its position in agriculture? J Pest Sci 93 (4): 1125–30.

Minh BQ, Schmidt HA, Chernomor O, Schrempf D, Woodhams MD, von Haeseler A, Lanfear R (2020) IQ-TREE 2: new models and efficient methods for phylogenetic inference in the genomic era. Mol Biol Evol 37 (5): 1530–34.

Mistry J, Finn RD, Eddy SR, Bateman A, Punta M (2013) Challenges in homology search: HMMER3 and convergent evolution of coiled-coil regions. Nucleic Acids Res 41 (12): e121.

Mistry J, et al. (2021) Pfam: The protein families database in 2021. Nucleic Acids Res 49 (D1): D412–19.

Mohr S, Bakal C, Perrimon N (2010) Genomic screening with RNAi: results and challenges. Annu Rev Biochem 79: 37–64.

Mohr SE, Smith JA, Shamu CE, Neumüller RA, Perrimon N (2014) RNAi screening comes of age: improved techniques and complementary approaches. Nat Rev Mol Cell Biol 15 (9): 591–600.

O’Callaghan M, Glare TR, Burgess EP, Malone LA (2005) Effects of plants genetically modified for insect resistance on nontarget organisms. Annu Rev Entomol 50: 271–92.

Onchuru TO, Kaltenpoth M (2019) Quantitative PCR primer design affects quantification of dsRNA-mediated gene knockdown. Ecol Evol 9 (14): 8187–92.

Paysan-Lafosse T, et al. (2023) InterPro in 2022. Nucleic Acids Res 51 (D1): D418–27.

Peng Y, et al. (2020) Identification of a double-stranded RNA-degrading nuclease influencing both ingestion and injection RNA interference efficiency in the red flour beetle *Tribolium castaneum*. Insect Biochem Mol Biol 125: 103440.

Peng Y, et al. (2021) Knockout of SldsRNase1 and SldsRNase2 revealed their function in dsRNA degradation and contribution to RNAi efficiency in the tobacco cutworm, Spodoptera litura. J Pest Sci 94 (4): 1449–60.

Perrimon N, Ni JQ, Perkins L (2010) In vivo RNAi: today and tomorrow. Cold Spring Harb Perspect Biol 2 (8): a003640.

Rakes M, Pasini RA, Morais MC, Araújo MB, de Bastos Pazini J, Seidel EJ, Bernardi D, Grützmacher AD (2021) Pesticide selectivity to the parasitoid *Trichogramma pretiosum*: a pattern 10-year database and its implications for Integrated Pest Management. Ecotoxicol Environ Saf 208: 111504.

Sharma R, Taning CNT, Smagghe G, Christiaens O (2021) Silencing of double-stranded ribonuclease improves oral RNAi efficacy in Southern green stinkbug *Nezara viridula*. Insects 12 (2): 115.

Smith SM (1996) Biological control with *Trichogramma*: advances, successes, and potential of their use. Annu Rev Entomol 41: 375–406.

Song H, et al. (2019) Contributions of dsRNases to differential RNAi efficiencies between the injection and oral delivery of dsRNA in *Locusta migratoria*. Pest Manag Sci 75 (6): 1707–17.

Vergani-Junior CA, Tonon-da-Silva G, Inan MD, Mori MA (2021) DICER: structure, function, and regulation. Biophysical Reviews 13 (6): 1081–90.

Wang M, Thomas N, Jin H (2017) Cross-kingdom RNA trafficking and environmental RNAi for powerful innovative pre- and post-harvest plant protection. Curr Opin Plant Biol 38: 133–41.

Wang ZZ, et al. (2018) Parasitic insect-derived miRNAs modulate host development. Nat Commun 9 (1): 2205.

Wilson RC, Doudna JA (2013) Molecular mechanisms of RNA interference. Annu Rev Biophys 42: 217–39.

Yan ZC, Hua HQ, Qi GY, Li YX (2022) Early detection and identification of parasitoid wasps *Trichogramma* Westwood (Hymenoptera: Trichogrammatidae) in their host eggs using polymerase chain reaction-restriction fragment length polymorphism. J Econ Entomol 115 (4): 1095–101.

Zang L, Wang S, Zhang F, Desneux N (2021) Biological control with *Trichogramma* in China: history, present status, and perspectives. Annu Rev Entomol 66: 463–84.

Zhang H, Li H-C, Miao X-X (2013) Feasibility, limitation and possible solutions of RNAi-based technology for insect pest control. Insect Sci 20 (1): 15–30.

Zhang X, Fan Z, Wang Q, Kong X, Liu F, Fang J, Zhang S, Zhang Z (2022) RNAi efficiency through dsRNA injection is enhanced by knockdown of dsRNA nucleases in the Fall Webworm, *Hyphantria cunea* (Lepidoptera: Arctiidae). Int J Mol Sci 23 (11).

