## Supplementary file for "Assessing RNAi feasibility and susceptibility to environmental RNAi in *Trichogramma dendrolimi* (Hymenoptera: Trichogrammatidae)"

**This supplemental file includes**

**Supplementary text**

**Supplementary tables**

**Table S1.** Primers used for dsRNA synthesis in this study

**Table S2.** Primers used for RT–qPCR in the study

**Table S3.** Characteristics of dsRNase genes in *T. dendrolimi* and *T. pretiosum*

**Table S4.** Additional target genes for environmental RNAi in *T. dendrolimi* adults and larvae

**Supplementary figures**

**Figure S1.** The maximum-likelihood phylogenetic tree of the RNase III family proteins.

**Figure S2.** Domain architecture of Dicer and Drosha proteins.

**Figure S3.** The maximum-likelihood phylogenetic tree of the Argonaute family proteins.

**Figure S4.** Domain architecture of Argonaute proteins.

**Figure S5.** The maximum-likelihood phylogenetic tree of the dsRNA-binding proteins.

**Figure S6.** Domain architecture of dsRNA-binding proteins.

**Figure S7.** The maximum-likelihood phylogenetic tree of the Sid-1 protein.

**Figure S8.** Domain architecture of Sid-1 proteins.

**Figure S9.** Lack of knockdown effects in *T. dendrolimi* adults following 3 days of continuous dsRNA feeding.

**Figure S10.** Lack of knockdown effects in *T. dendrolimi* larvae parasitized in artificial hosts containing dsRNA.

### Supplementary Text

#### Phylogenetic and domain analysis of *Trichogramma* genes in the RNAi pathway

#### (1) RNaseIII

In the RNAi pathways of insects, there are three proteins belonging to the RNase III family, Dicer-1, Dicer-2, and Drosha (Carthew and Sontheimer 2009; Vergani-Junior et al. 2021; Wilson and Doudna 2013). Two Dicers and one Drosha were identified in the genomes of both *T. dendrolimi* and *T. pretiosum*. The identified *Trichgoramma* proteins were subjected to phylogenetic analysis with the RNase III family proteins from *Apis mellifera* (Dicer-1: NP_001116485.2, Dicer-2: XP_016773223.2, Drosha: XP_016766928.1), *Drosophila melanogaster* (Dicer-1: NP_524453.1, Dicer-2: NP_523778.2, Drosha: NP_477436.1), *Tribolium castaneum* (Dicer-1: EFA11550.2, Dicer-2: NP_001107840.1, Drosha: XP_008199088.1), and *Bombyx mori* (Dicer-1: XP_037869731.1, Dicer-2: NP_001180543.1, Drosha: XP_012547603.2). The results showed that the RNase III proteins were divided into two clades, with Dicer-1 and Dicer-2 proteins clustering together in one clade, while Drosha proteins formed a separate clade (Fig. S1). Dicer1, Dicer2, and Drosha proteins from *Trichogramma* have the closest orthologous relationships with those from *Apis mellifera* (Fig. S1). Domain analysis revealed that the domain composition and arrangement in Dicer-1 were consistent, including Helicase_C (PF00271.34), Dicer_dimer (PF03368.17), PAZ (PF02170.25), and Ribonuclease_3 (PF00636.29) (Fig. S2a). Dicer2 proteins contained similar domains but with an additional DEAD (PF00270.32) domain at the N-terminus, and Dicer2 proteins from *A. mellifera* and *D. melanogaster* also contained a dsrm (PF00035.29) domain at the C-terminus (Fig. S2b). All Drosha proteins contain only Ribonuclease_3 and dsrm domains (Fig. S2c). The domain composition and arrangement of *Trichogramma* Dicer-1, Dicer-2 and Drosha are similar to those from other insects.

#### (2) Argonaute

Argonaute proteins are essential components of the RNA-induced silencing complex (RISC), which is responsible for binding and cleave the target RNA, leading to its degradation or inhibition of translation (Carthew and Sontheimer 2009; Meister 2013; Wilson and Doudna 2013). Three Argonaute proteins were identified in the genomes of both *T. dendrolimi* and *T. pretiosum*. The identified *Trichgoramma* proteins were subjected to phylogenetic analysis with the Argonaute family proteins from *A. mellifera* (Ago-1: XP_006571833.1, Ago-2: XP_395048.4), *D. melanogaster* (Ago-1: NP_001246314.1, Ago-2: NP_648775.1), *T. castaneum* (Ago-1: XP_008196653.1, Ago-2a: NP_001107842.1), and *B. mori* (Ago-1: NP_001095931.1, Ago-2: NP_001036995.2). The results showed that Ago-1 and Ago-2 proteins are divided into two separate clades (Fig. S3). In the clade of Ago-1, *Trichogramma* Ago-1a and Ago-1b proteins were clustered together with AmelAgo-1 as the closest outgroup, suggesting a recent gene duplication of Ago-1 in *Trichogramma* after its divergence from *A. mellifera* (Fig. S3). Domain analysis revealed that the domain composition and arrangement of Ago-1 and Ago-2 are similar, including ArgoN (PF16486.8), ArgoL1 (PF08699.13), PAZ (PF02170.25), ArgoL2 (PF16488.8), ArgoMid (PF16487.8), and Piwi (PF02171.20) (Fig. S4). However, the ArgoMid domain is generally absent in Ago-2 proteins, except in TcasAgo-2a and DmelAgo-2 (Fig. S4b). The domain composition and arrangement of *Trichogramma* Argonaute proteins are similar to those from other insects.

**(3) dsRNA binding protein**

In the RNAi pathway of insects, dsRNA-binding proteins generally function as cofactors for RNase III proteins (Carthew and Sontheimer 2009; Wilson and Doudna 2013). One Lops, one Pasha, and one R2D2 proteins were identified in the genomes of both *T. dendrolimi* and *T. pretiosum*. The identified *Trichogramma* proteins were subjected to phylogenetic analysis with dsRNA-binding proteins from *A. mellifera* (Lops: KAG6801533.1, Pasha: XP_006559676.1, R2D2: XP_006560091.1), *D. melanogaster* (Lops: NP_723813.1, Pasha: NP_001263149.1, R2D2: NP_609152.1), *T. castaneum* (Lops: XP_008198428.1, Pasha: XP_015836202.1, R2D2: NP_001128425.1), *B. mori* (Lops: XP_021207536.2, Pasha: XP_037872173.1, R2D2: NP_001182007.1). The results showed that Lops and R2D2 had a closer phylogenetic relationship, while their relationship with Pasha was more distant (Fig. S5). Among these dsRNA-binding proteins, *Trichogramma* proteins have the closest orthologous relationships with those from *A. mellifera* (Fig. S5). Domain analysis revealed that dsRNA-binding proteins contain dsrm (PF00035.29) domains (Fig. S6). Both Lops and R2D2 proteins contained two dsrm domains, while Pasha proteins had one dsrm domain (Fig. S6). The domain composition and arrangement in *Trichogramma* proteins are similar to those from other insects.

**(4) Transmembrane channel protein**

Although the Sid-1 proteins do not directly participate in the RNAi pathway, their ability to transmit RNAi signals has been discovered in various organisms (Carthew and Sontheimer 2009; Wilson and Doudna 2013). We also analyzed Sid-1 and identified two Sid-1 proteins in the genomes of both *T. dendrolimi* and *T. pretiosum*. These identified *Trichgoramma* proteins were subjected to phylogenetic analysis with the Sid-1 proteins from *A. mellifera* (sid-1: XP_006565236.1), *B. mori* (sid-1: NP_001106735.1), *Helicoverpa armigera* (sid-1: XP_049707213.*1), Hyphantria cunea* (sid-1: AXZ00255.1), *Leptinotarsa decemlineata* (Sid-1: ALG36906.1), *Locusta migratoria manilensis* (Sid-1: AFQ00936.1), *Spodoptera litura* (Sid-1: AHC98014.1), and *T. castaneum* (Sid-1a: ABU63672.1, Sid-1b: ABU63673.1, sid-1c: ABU63674.1). The results showed that the *Trichogramma* sid-1 proteins form a clade with AmelSid-1 (Fig. S7). Domain analysis revealed that Sid-1 proteins all contain only one SID-1_RNA_chan (PF13965.9) domain (Fig. S8). The SID-1_RNA_chan domains in different Sid-1 proteins have similar lengths, except for the shorter domain length in TdenSid-1b compared to other insects (Fig. S8).

**Table S1. Primers used for dsRNA synthesis in this study**

| Primer | Sequence |
|---|---|
| CHT10-FP | TAATACGACTCACTATAGGGATGACCTGAAAATAATCGC |
| CHT10-RP | TAATACGACTCACTATAGGGGAAATAAGCCAGCTCTCC |
| VAA-FP | TAATACGACTCACTATAGGGGAGGAGAGGGAGGCTAAGTT |
| VAA-RP | TAATACGACTCACTATAGGGTTGGTGTGTGCGAGAAAGA |
| VAB-FP | TAATACGACTCACTATAGGGGGAGACCGCTCGTTTCTT |
| VAB-RP | TAATACGACTCACTATAGGGTGTCGTCGTTGGGCATAGT |
| G3PDH-FP | TAATACGACTCACTATAGGGGCCAAATGTCATCGGTAT |
| G3PDH-RP | TAATACGACTCACTATAGGGGCAGCTCTCGAAGAAGGT |
| IAP-FP | TAATACGACTCACTATAGGGGGCAATCCTTTCCTGTCT |
| IAP-RP | TAATACGACTCACTATAGGGGTGCTAGTGCTGGTGGGT |
| shrub-FP | TAATACGACTCACTATAGGGTCAGTGGGAAGAAAGAGG |
| shrub-RP | TAATACGACTCACTATAGGGGACCAAAGGCTACAGGAT |
| HSC-FP | TAATACGACTCACTATAGGGCACTACGCCCAGTTATGT |
| HSC-RP | TAATACGACTCACTATAGGGGCTGCGAGTCGTTGAAAT |
| MAS-FP | TAATACGACTCACTATAGGGGCTCTGGTCTGAAGGGTA |
| MAS-RP | TAATACGACTCACTATAGGGCTTCTGACCGATAGCGAC |
| ENO-FP | TAATACGACTCACTATAGGGGAAACTTGACGGAACACC |
| ENO-RP | TAATACGACTCACTATAGGGTTACCTTCATAACCAGCAG |
| SPI-FP | TAATACGACTCACTATAGGGGGCTTGACAATGGCAGAC |
| SPI-RP | TAATACGACTCACTATAGGGGAGAAGAAGTTGGTGGCG |
| VAE-FP | TAATACGACTCACTATAGGGTTAGTACAGCAACAGCGT |
| VAE-RP | TAATACGACTCACTATAGGGAGCAGCCAAGTAGTTATCT |

### Table S2. Primers used for RT–qPCR in the study

| Prime | Sequence |
|---|---|
| qCHT10-FP | CAACAAGCACGACAGACCTATTC |
| qCHT10-RP | GGGGCTACGGGTTTTTCAC |
| qVAA-FP | GGAAACAGAATTCGACGGA |
| qVAA-RP | TACACAGGGGAACAGGGAG |
| qVAB-FP | TCCAGGGTCAACCCATCA |
| qVAB-RP | AAACCAGCAGCCGAGAAA |
| qIAP-FP | ATTCATCAACAGCGTCACCG |
| qIAP-RP | GGGTTTCCTCTTTGCTTTCAC |
| qshrub-FP | ATTTGCCTGCTGAGCCTGTT |
| qshrub-RP | ACGCTGCCCATTGTTCCA |
| qHSC-FP | CACCTTTGATGTTTCCATTCTG |
| qHSC-RP | GGTTGTCGAAGTCTTCCCC |
| qATPase-FP | CTTCAGAGATAACGGCAAACA |
| qATPase-RP | AGGAGACGAGAGTGGAGGTAG |
| qENO-FP | ACTTCAAGAACCCCAACTCC |
| qENO-RP | TAGTCCACGAATCCCAGCC |
| qSPI-FP | TCGTCTGGCAAAGAGCGGT |
| qSPI-RP | TGAAATCGGCAGCGGGACT |
| qVAE-FP | TGGTGTTGAACTTTTGGCCC |
| qVAE-RP | CGTTGCGACCGAATAATGC |

**Table S3. Characteristics of dsRNase genes in *T. dendrolimi* and *T. pretiosum***

| Gene | Length (aa.) | IP | MW (kDa) | Accession number |
|---|---|---|---|---|
| ***T. dendrolimi*** | | | | |
| endoG | 235 | 9.15 | 27.15 | Tden02749.t1 |
| dsRNase-1 | 536 | 8.29 | 61.23 | Tden04820.t1 |
| dsRNase-2 | 467 | 8.92 | 53.25 | Tden05237.t1 |
| dsRNase-3 | 447 | 8.38 | 50.13 | Tden06522.t1 |
| dsRNase-4 | 445 | 7.21 | 50.13 | Tden06529.t1 |
| dsRNase-5 | 481 | 6.00 | 54.49 | Tden06530.t1 |
| dsRNase-6 | 465 | 8.80 | 53.34 | Tden06663.t1 |
| dsRNase-7 | 272 | 9.54 | 31.59 | Tden06664.t1 |
| dsRNase-8 | 461 | 8.95 | 53.24 | Tden06665.t1 |
| dsRNase-9 | 582 | 9.77 | 66.24 | Tden06779.t1 |
| dsRNase-10 | 509 | 9.18 | 57.37 | Tden06780.t1 |
| dsRNase-11 | 561 | 9.29 | 63.42 | Tden06784.t1 |
| dsRNase-12 | 730 | 8.80 | 86.67 | Tden07233.t1 |
| dsRNase-13 | 481 | 5.43 | 54.98 | Tden07665.t1 |
| dsRNase-14 | 780 | 8.24 | 87.35 | Tden07667.t1 |
| dsRNase-15 | 402 | 8.93 | 44.98 | Tden11671.t1 |
| dsRNase-16 | 803 | 8.95 | 95.40 | Tden15507.t1 |
| ***T. pretiosum*** | | | | |
| endoG | 321 | 9.27 | 36.33 | XP_014233302.1 |
| dsRNase-1 | 429 | 9.08 | 48.27 | XP_014221399.1 |
| dsRNase-2 | 491 | 5.78 | 55.29 | XP_014226211.1 |
| dsRNase-3 | 468 | 8.19 | 52.10 | XP_014226213.1 |
| dsRNase-4 | 466 | 8.90 | 53.32 | XP_014233059.1 |
| dsRNase-5 | 499 | 9.24 | 56.12 | XP_014233529.2 |
| dsRNase-6 | 480 | 5.51 | 54.77 | XP_023313547.1 |
| dsRNase-7 | 506 | 8.73 | 57.21 | XP_023314185.1 |
| dsRNase-8 | 402 | 8.85 | 44.96 | XP_023314495.1 |
| dsRNase-9 | 430 | 9.32 | 48.99 | XP_023317907.1 |
| dsRNase-10 | 191 | 6.37 | 22.08 | XP_023318433.1 |
| dsRNase-11 | 468 | 8.98 | 53.35 | XP_023319337.1 |

aa. : amino acids; IP: isoelectric point; MW: molecular weight.

**Table S4. Additional target genes for environmental RNAi in *T. dendrolimi* adults and larvae**

| Abbreviation | Gene name | ORF (bp) | Length (aa.) | Accession number |
|---|---|---|---|---|
| G3PHD | *Glyceraldehyde-3-phosphate dehydrogenase* | 1111 | 370 | Tden06713.t1 |
| IAP | *Inhibitor of apoptosis protein* | 1378 | 459 | Tden14259.t1 |
| *Shrb* | *Shrb* | 664 | 221 | Tden09551.t1 |
| HSC | *Heat shock protein* | 1969 | 656 | Tden01651.t1 |
| MAS | *ATP synthase subunit alpha* | 1642 | 547 | Tden13395.t1 |
| SPI | *Serine protease inhibitor* | 1198 | 399 | Tden11342.t1 |
| VAE | *v-ATPase E* | 682 | 227 | Tden12157.t1 |
| ENO | *Enolase* | 1381 | 460 | Tden14855.t1 |

ORF: open reading frame; aa. : amino acids.

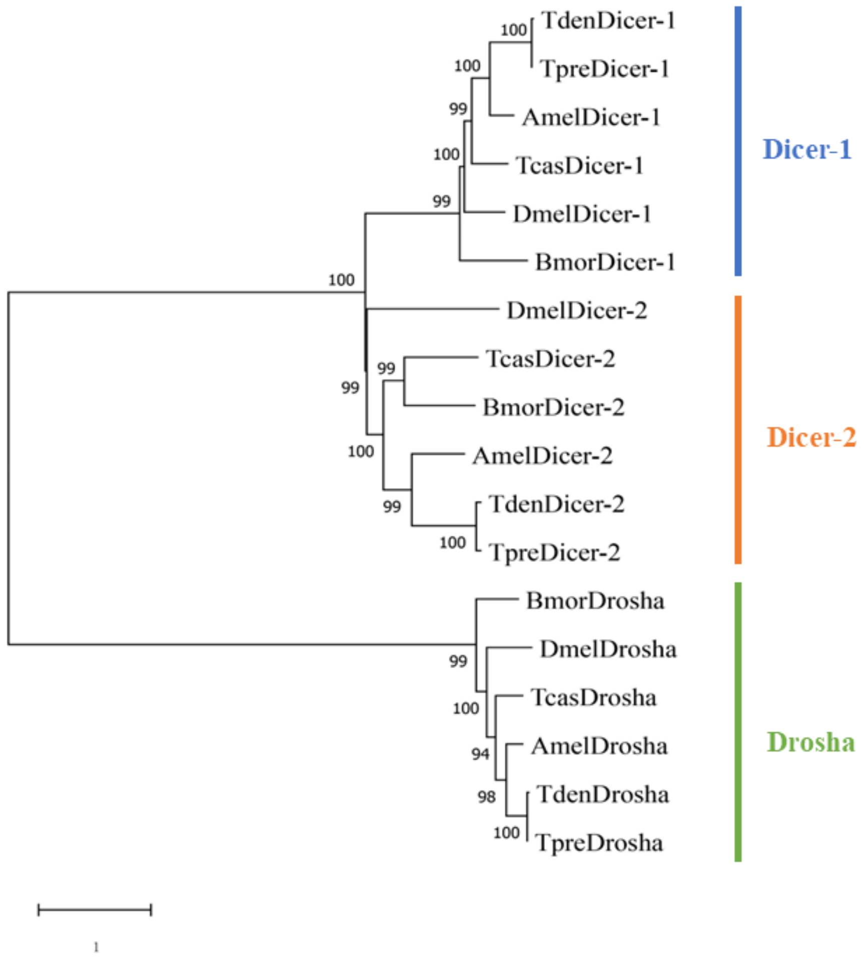


**Figure S1. The maximum-likelihood phylogenetic tree of the RNase III family proteins.** Amel: *Apis mellifera*; Bmor: *Bombyx mori*; Dmel: *Drosophila melanogaster*; Tcas: *Tribolium castaneum*; Tden: *Trichogramma dendrolimi*; Tpre: *Trichogramma pretiosum*.

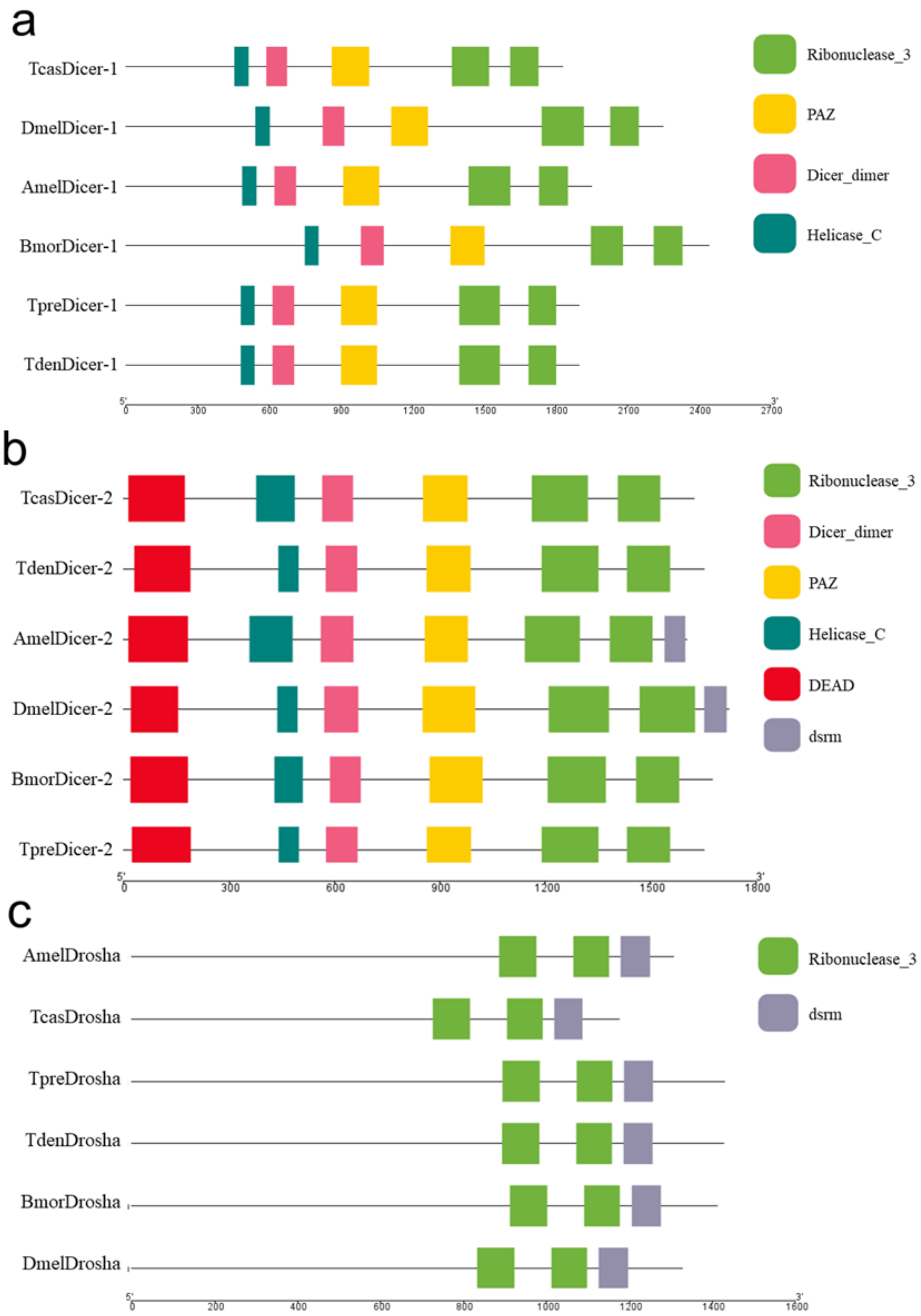


**Figure S2. Domain architecture of Dicer and Drosha proteins.** The protein sequences of (a) Dicer-1, (b) Dicer-2 and (c) Drosha were annotated by the Pfam and Interpro databases, and the domain information was visualized by TBtools. Amel: *Apis mellifera*; Bmor: *Bombyx mori*; Dmel: *Drosophila melanogaster*; Tcas: *Tribolium castaneum*; Tden: *Trichogramma dendrolimi*; Tpre: *Trichogramma pretiosum*.

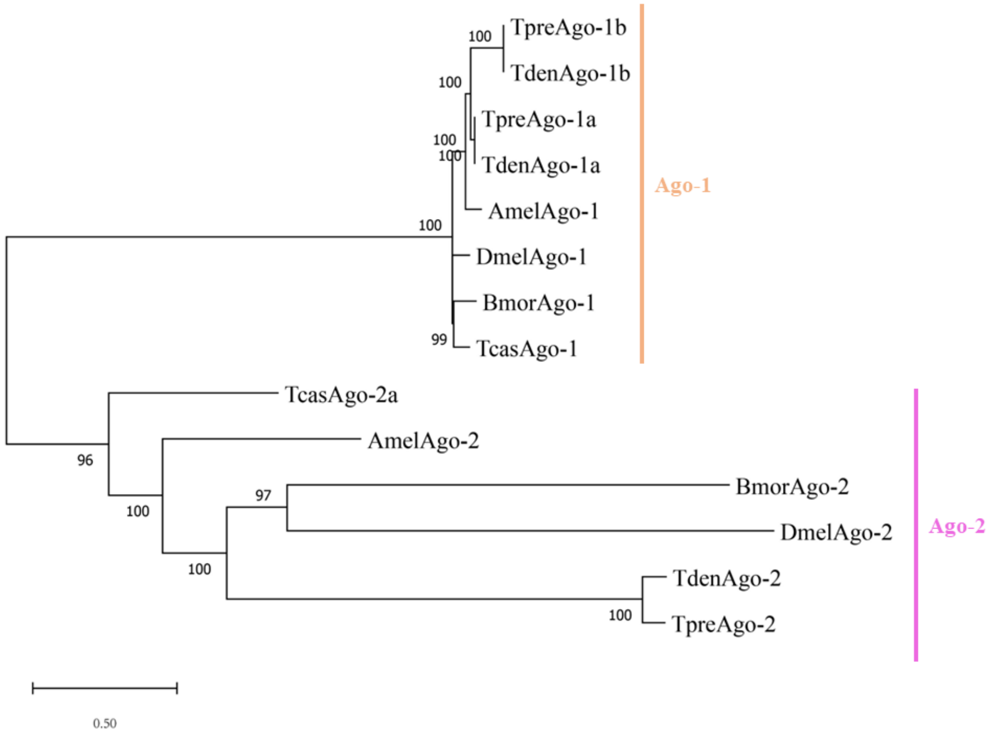


**Figure S3. The maximum-likelihood phylogenetic tree of the Argonaute family proteins.** Amel: *Apis mellifera*; Bmor: *Bombyx mori*; Dmel: *Drosophila melanogaster*; Tcas: *Tribolium castaneum*; Tden: *Trichogramma dendrolimi*; Tpre: *Trichogramma pretiosum*.

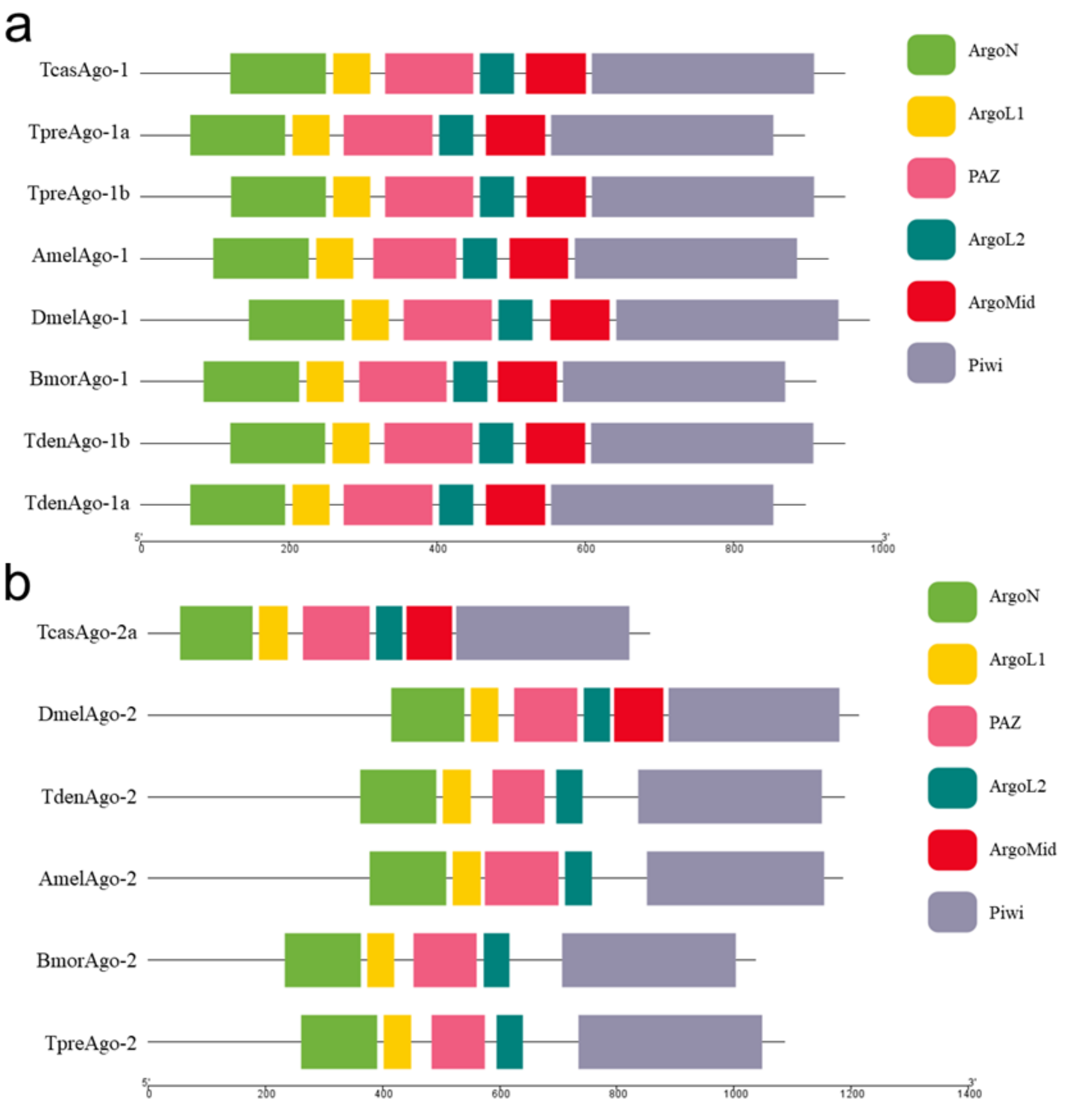


**Figure S4. Domain architecture of Dicer and Drosha proteins.** The protein sequences of (b) Ago-1 and (c) Ago-2 were annotated by the Pfam and Interpro databases, and the domain information was visualized by TBtools. Amel: *Apis mellifera*; Bmor: *Bombyx mori*; Dmel: *Drosophila melanogaster*; Tcas: *Tribolium castaneum*; Tden: *Trichogramma dendrolimi*; Tpre: *Trichogramma pretiosum*.

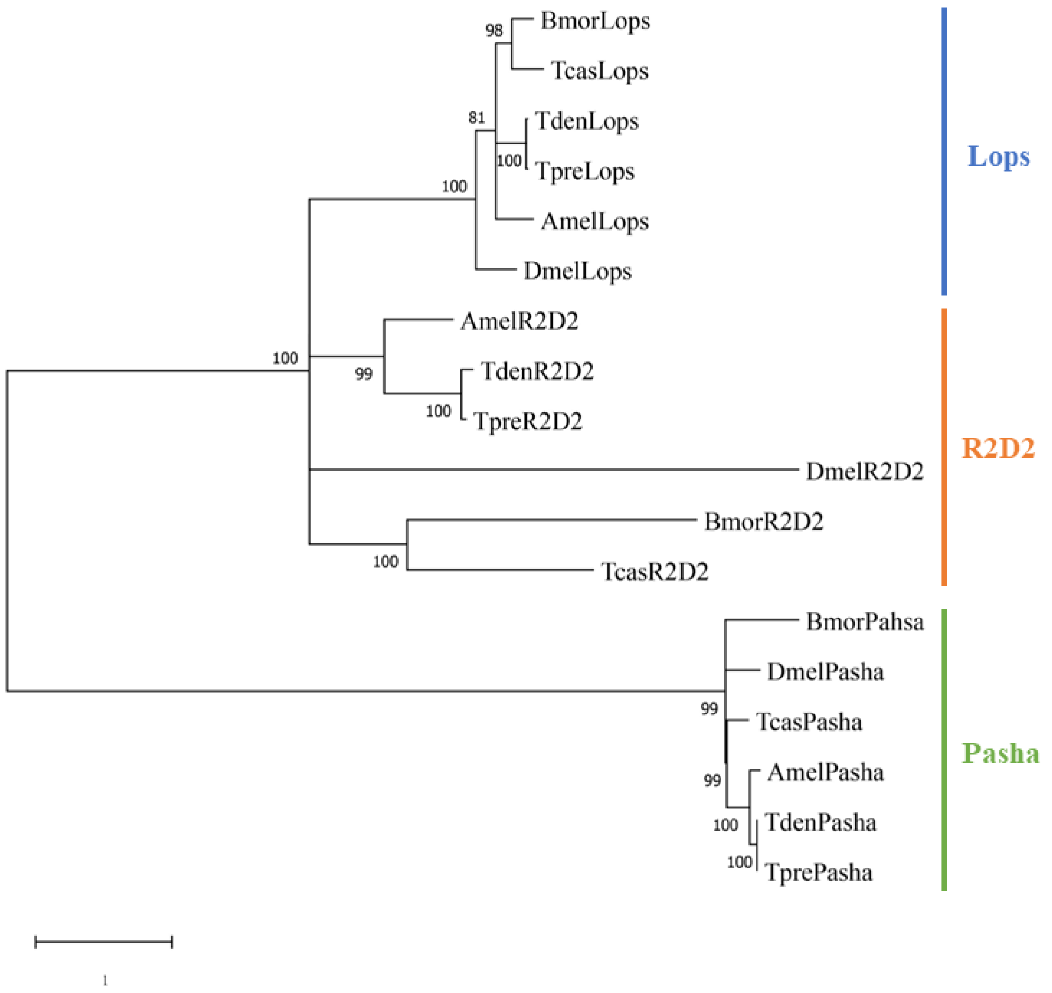


**Figure S5. The maximum-likelihood phylogenetic tree of the dsRNA-binding proteins.** Amel: *Apis mellifera*; Bmor: *Bombyx mori*; Dmel: *Drosophila melanogaster*; Tcas: *Tribolium castaneum*; Tden: *Trichogramma dendrolimi*; Tpre: *Trichogramma pretiosum*.

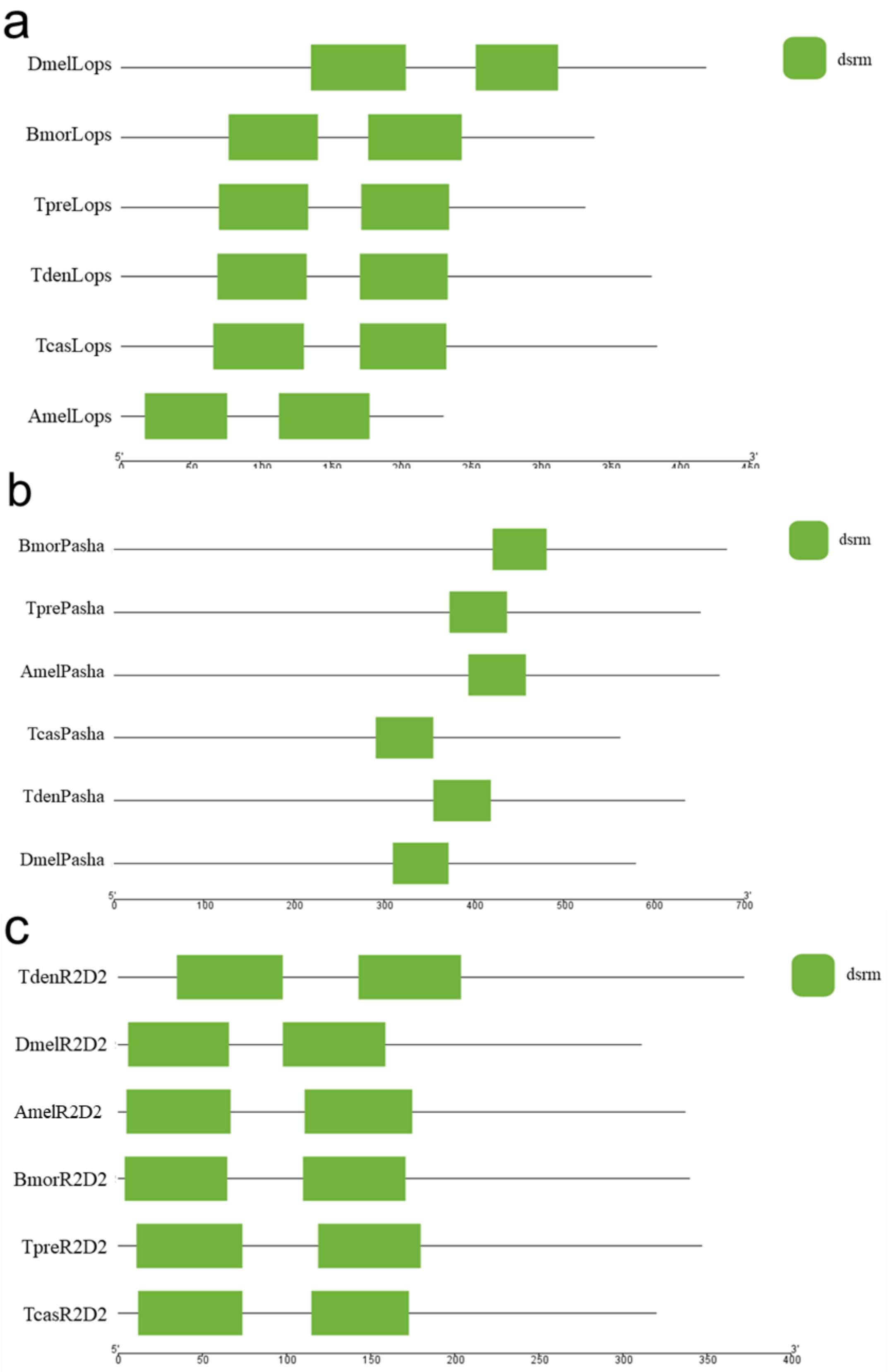


**Figure S6. Domain architecture of Dicer and Drosha proteins.** The protein sequences of (a) Lops, (b) Pasha and (c) R2D2 were annotated by the Pfam and Interpro databases, and the domain information was visualized by TBtools. Amel: *Apis mellifera*; Bmor: *Bombyx mori*; Dmel: *Drosophila melanogaster*; Tcas: *Tribolium castaneum*; Tden: *Trichogramma dendrolimi*; Tpre: *Trichogramma pretiosum*.

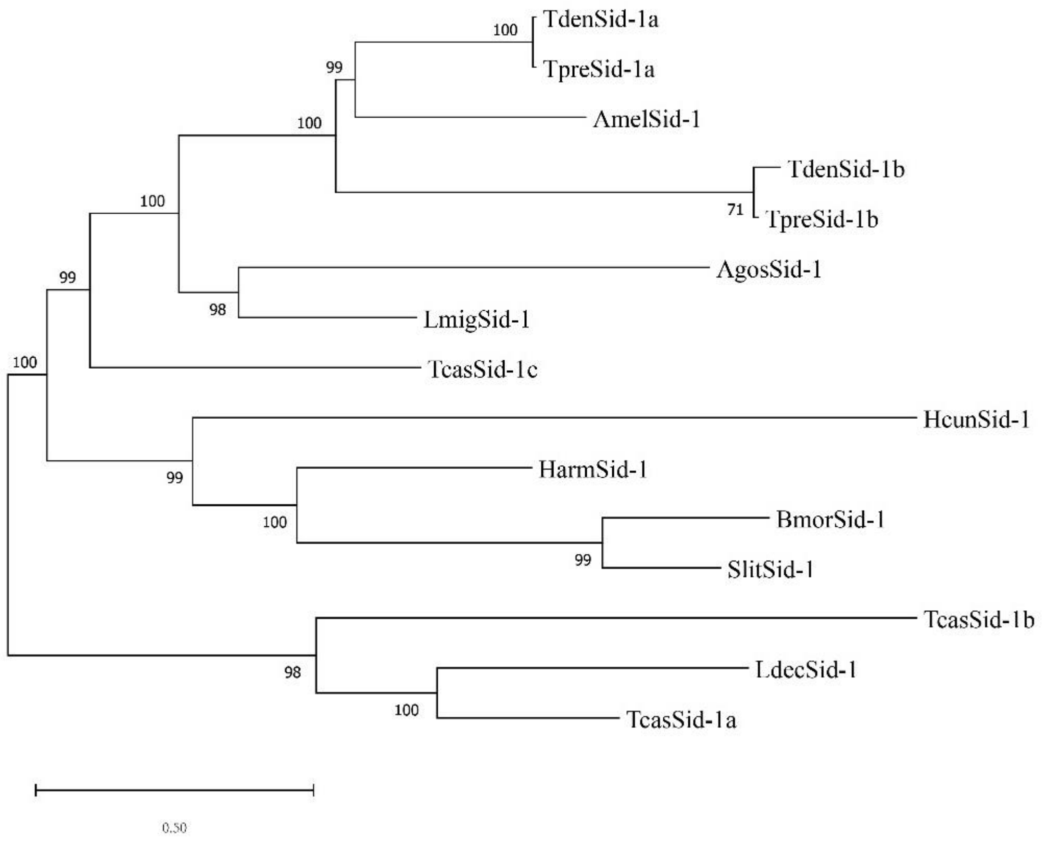


**Figure S7. The maximum-likelihood phylogenetic tree of the Sid-1 protein.**

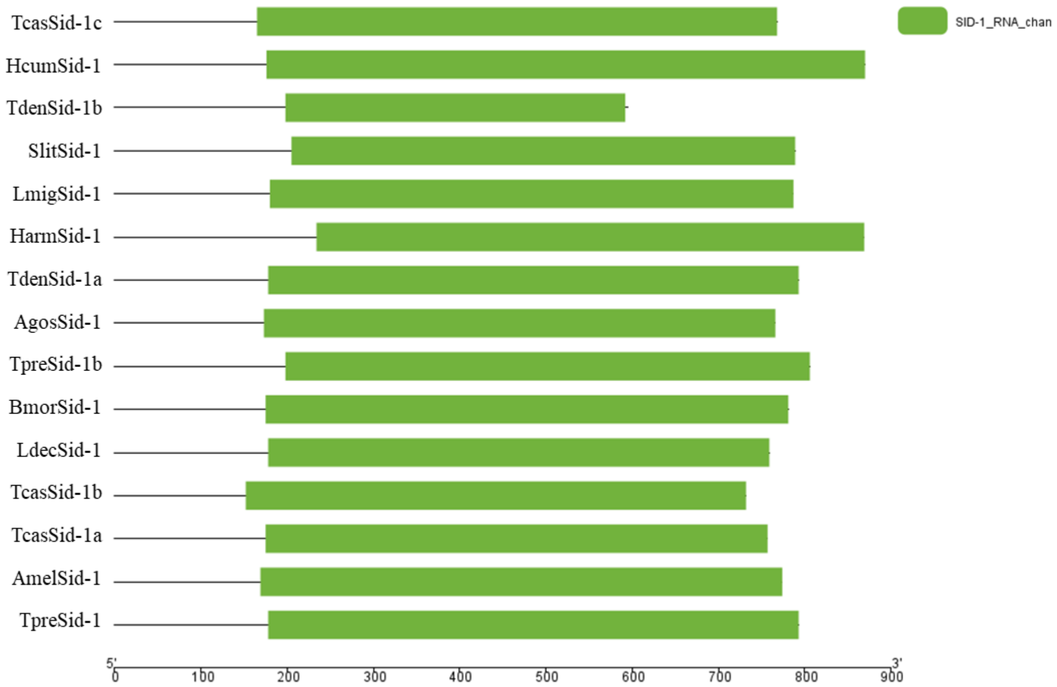


**Figure S8. Domain architecture of Sid-1 proteins.** Amel: *Apis mellifera*; Bmor: *Bombyx mori*; Dmel: *Drosophila melanogaster*; Harm: *Helicoverpa armigera*; Hcun: *Hyphantria cunea*; Ldec: *Leptinotarsa decemlineata*; Lmig: *Locusta migratoria manilensis*; Slit: *Spodoptera litura*; Tcas: *Tribolium castaneum*; Tden: *Trichogramma dendrolimi*; Tpre: *Trichogramma pretiosum*.

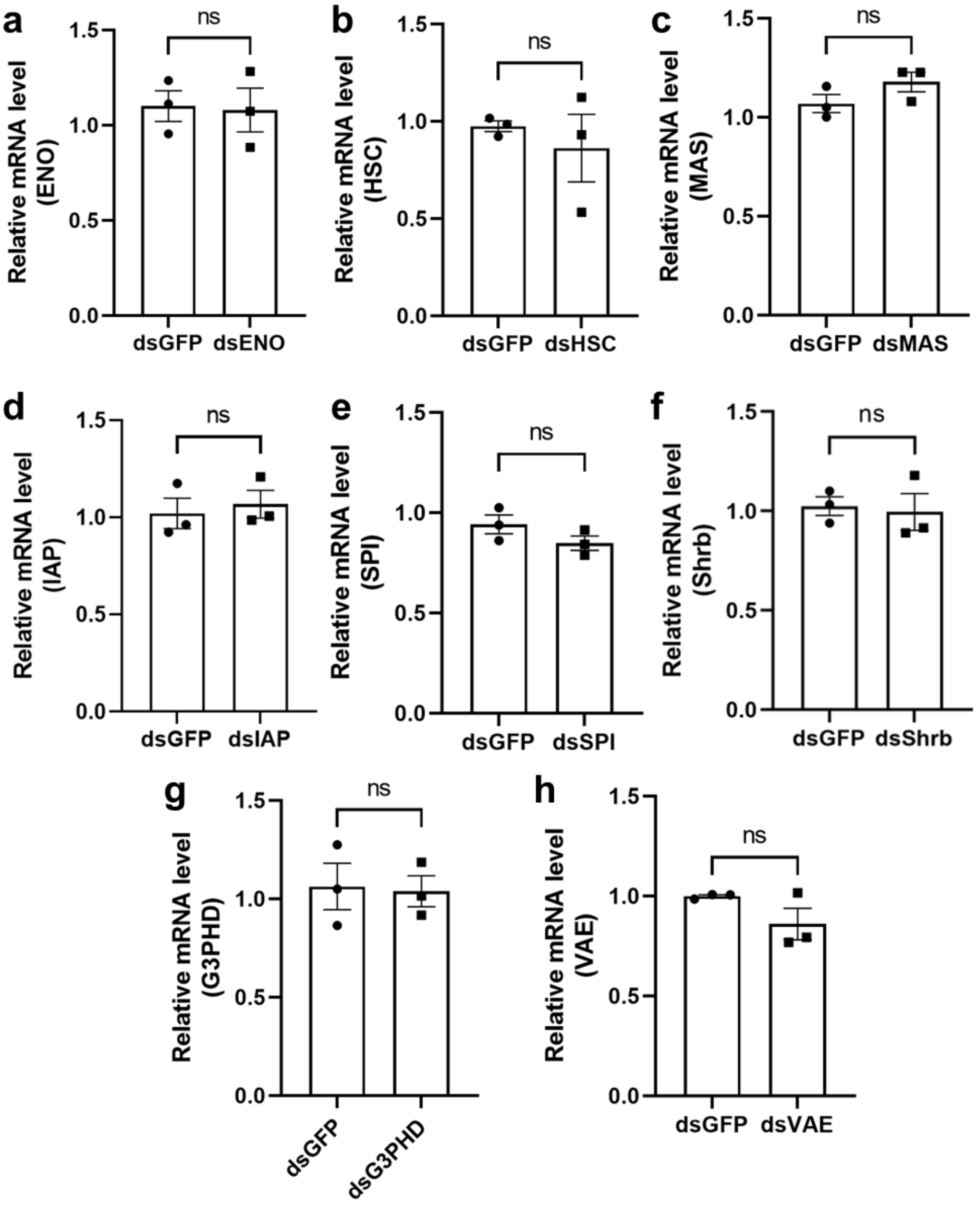


**Figure S9. Lack of knockdown effects in *T. dendrolimi* adults following 3 days of continuous dsRNA feeding.** Relative expression levels of (a) ENO, (b) HSC, (c) MAS, (d) IAP, (e) SPI, (f) Shrb, (g) G3PHD, and (h) VAE in *T. dendrolimi* adults after feeding on dsRNA. GFP dsRNA served as a negative control. Data are displayed as the mean ± SEM, with n=3. Ns: no significant difference.

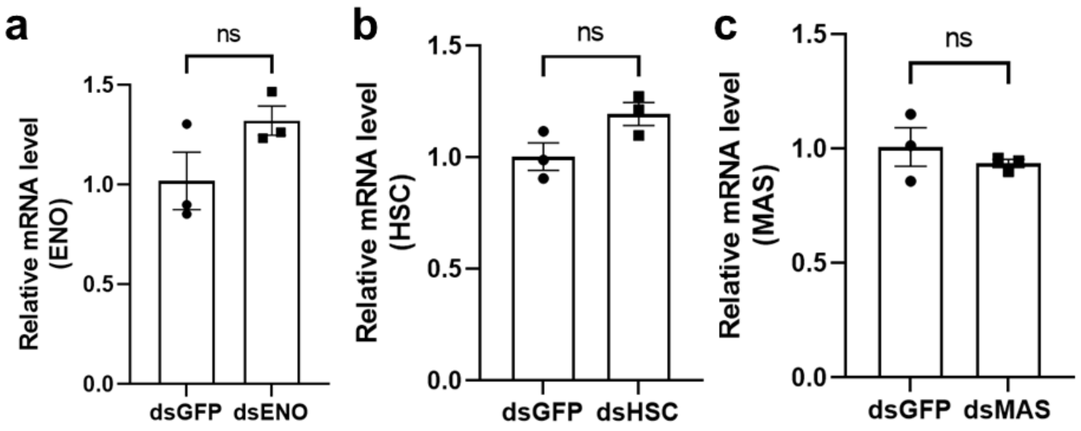


**Figure S10. Lack of knockdown effects in *T. dendrolimi* larvae parasitized in artificial hosts containing dsRNA.** Relative expression levels of (a) ENO, (b) HSC, and (c) MAS in *T. dendrolimi* larvae 3 days after parasitism in artificial hosts containing dsRNA. GFP dsRNA served as a negative control. Data are displayed as the mean ± SEM, with n=3. Ns: no significant difference.
